# Evolutionary and spike-timing-dependent reinforcement learning train spiking neuronal network motor control

**DOI:** 10.1101/2021.11.20.469405

**Authors:** Daniel Haşegan, Matt Deible, Christopher Earl, David D’Onofrio, Hananel Hazan, Haroon Anwar, Samuel A Neymotin

## Abstract

Despite being biologically unrealistic, artificial neural networks (ANNs) have been successfully trained to perform a wide range of sensory-motor behaviors. In contrast, the performance of more biologically realistic spiking neuronal network (SNN) models trained to perform similar behaviors remains relatively suboptimal. In this work, we aimed at pushing the field of SNNs forward by exploring the potential of different learning mechanisms to achieve optimal performance. Inspired by biological learning mechanisms operating at multiple timescales, we used spike-timing-dependent reinforcement learning (STDP-RL) and evolutionary strategy (EVOL) with SNNs to solve the CartPole reinforcement learning (RL) control problem. Though the role of STDP-RL in biological systems is well established, several other mechanisms, though not fully understood, work in concert during learning in vivo. Recreating accurate models that capture the interaction of STDP-RL with these diverse learning mechanisms is extremely difficult. EVOL is an alternative method, and has been successfully used in many studies to fit model neural responsiveness to electrophysiological recordings and in some cases for classification problems. One advantage of EVOL is that it may not need to capture all interacting components of synaptic plasticity, and thus provides a better alternative to STDP-RL. Here, we compared the performance of each algorithm after training, which revealed EVOL as a powerful method to training SNNs to perform sensory-motor behaviors. Our modeling opens up new capabilities for SNNs in RL and could serve as a testbed for neurobiologists aiming to understand multi-timescale learning mechanisms and dynamics in neuronal circuits.

## Introduction

Reinforcement Learning (RL) problems offer an ideal framework for comparing learning strategies of an interactive, goal-seeking agent (Sutton and Barto 2018). Most often, the best learning strategy is evaluated based on the time efficiency and algorithm complexity. While there are many deep reinforcement learning algorithms for solving dynamical control problems (Mnih et al. 2015), biologically realistic network architectures and training strategies are not yet as efficient. In this work, using the CartPole RL problem, we compare biologically inspired learning algorithms based on the training efficiency and the resulting network dynamics.

As Spiking Neural Networks (SNNs) are shown to be Turing-complete (Maass 1996b), and computationally more powerful than Artificial Neural Networks (ANNs) (Maass 1997, [a] 1996), efficient learning strategies are still actively investigated (Tavanaei et al. 2019). SNNs have been effective for pattern recognition problems (Gupta and Long 2007; Escobar et al. 2009; Kasabov et al. 2014; Tavanaei and Maida 2017; Mozafari et al. 2018) but are rarely used for solving reinforcement learning control problems. As spiking neurons operate in the time domain, we show that RL problems are suitable for evaluating training strategies and for providing insight into neural circuit dynamics.

Traditionally, when SNN models are trained to perform a behavior using biologically inspired learning mechanisms, algorithms used are variations on either Spike Timing Dependent Plasticity(STDP) (Tavanaei et al. 2019) or Evolutionary Strategies (Espinal et al. 2014). For learning behaviors from the reinforcement learning domain, STDP can be extended to use reward modulated plasticity (Anwar et al. 2021; Patel et al. 2019; Hazan et al. 2018; Chadderdon et al. 2012; Neymotin et al. 2013), an algorithm denoted Spike-timing dependent reinforcement learning (STDP-RL). In this work, we introduce a new Evolutionary Strategy variation, adapted from non-spiking neural networks (Salimans et al. 2017). Alternatively, there are many algorithms that learn behaviors through backpropagation(Bohte et al., 2002, Liu et al., 2017, Mostafa 2017, Wu et al., 2017, Huh et al., 2017). Since the biological plausibility of backpropagation is still debated (Stork 1989; Mazzoni, Andersen, and Jordan 1991; Whittington and Bogacz 2019) we have not used it in this work.

STDP-RL trains SNNs by establishing associations between the neurons encoding the sensory environment and neurons producing an action or sequence of actions, such that appropriate actions are produced for specific sensory cues. The sensory-motor associations are established from reward-modulated synaptic weight changes that occur at each timestep of the simulation. Hence, STDP-RL trains at the individual level, as we consider each separate initialization of an SNN network a separate “individual”.

In our SNNs, we simulate individual neurons as event-based dynamical units that mimic functions of their biological counterparts, like adaptation, bursting, and depolarization blockade (Neymotin et al. 2011). For training SNNs based on population-level fitness metrics, we adapt Evolutionary Strategies (EVOL) (Salimans et al. 2017) that has been shown to be efficient in training Artificial Neural Networks on similar problems (Salimans et al. 2017; Chrabaszcz, Loshchilov, and Hutter 2018). The EVOL algorithm treats the synaptic weight as an individual’s genome, then perturbs the genome using the mutation genetic operator to produce an offspring population. Each offspring is an individual SNN that interacts with the environment by receiving sensory stimuli and producing motor actions based on its internal firing patterns. Based on the offspring fitness, we combine the population genomes to yield the next generation of genomes. In the EVOL algorithm, SNN weights are not adapting throughout the interaction with the environment.

To summarize, in this work, we investigate two biologically inspired algorithms for training SNNs on the CartPole RL problem: individual-level training using STDP-RL, and population-level Evolutionary Strategies (EVOL). Our contributions are as follows: (1) we analyze the efficiency of biologically inspired training strategies for solving the CartPole RL problem; (2) we analyze the effect between individual-level and population-level learning on SNNs sensorimotor mappings and neuronal dynamics.

## Materials and Methods

### CartPole Game

To test different learning strategies, we chose the classic CartPole problem of balancing a vertical pole on a cart (Barto, Sutton, and Anderson 1983; Geva and Sitte 1993). All the simulations were run using the CartPole-v1 environment (**Figure 1 left**) available in the OpenAI Gym platform (Brockman et al. 2016) (https://gym.openai.com). To keep the pole balanced (**Figure 1 left**), a force to the left (−1) or the right (+1) must be applied at each time step of the simulation. Once the force is applied, a new game-state is generated, resulting from the interaction between the previous game-state and the applied action. The environment is fully described by four observations: cart position, cart horizontal velocity, pole angle with respect to a vertical axis, and angular velocity of the pole. For simplicity, we will now reference those four observations as position, velocity, angle, and angular velocity (see **Figure 1 left**). The position is a relative distance from the center of the screen with positive (negative) values to the right (left). Similarly, the pole’s angle represents positive values for angles to the right (clockwise). The velocity and angular velocity represents the rate of change of the position and the angle, respectively. The game is played in episodes, where each episode consists of multiple steps (left or right) taken to keep the pole balanced. The pole loses its balance when either the angle is greater than 15 degrees from vertical in either direction or the position is more than 2.4 units from the center in either direction. Each episode is allowed a maximum duration of 500 steps. An episode can be instantiated to different initial positions, which deterministically affects the trajectory through observational space.

**Figure 1:**
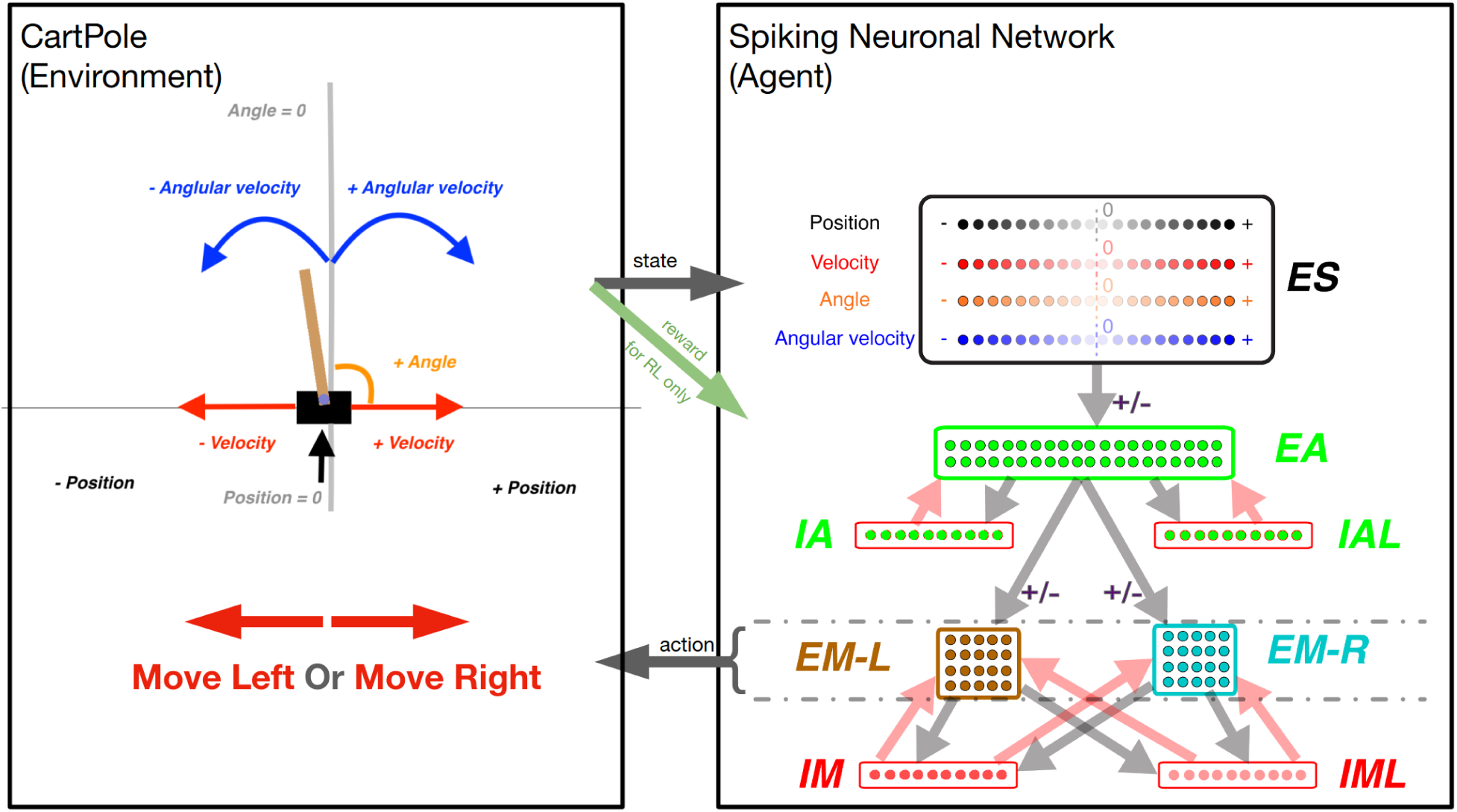
The CartPole game environment (left) interfacing with the SNN model (right). **Left box**: At each game-step, a new game state is produced which is described by the game state variables (as labeled): Position, Velocity, Angle, and Angular Velocity. **Arrow “state”**: The values of these variables are used to activate a unique quadruple of neurons in the “ES” neuronal population (1 for each state variable). **Right box:** The neuronal network is represented as a diagram, as each dot represents one neuron, and arrows (light gray and light red) represent connectivity between populations of neurons. The light gray and light red arrows represent excitatory and inhibitory synapses between neuronal populations respectively. There are four excitatory neuronal populations: “ES” with 80 neurons (black-bordered box); “EA” with 40 neurons (green-bordered box); “EM-L” with 20 neurons (brown-bordered box); and “EM-R” with 20 neurons (cyan-bordered box). There are four inhibitory neuronal populations: “IA” with 10 neurons; “IAL” with 10 neurons; “IM” with 10 neurons; and “IML” with 10 neurons (each within a red-bordered box). Excitatory neuronal populations have outgoing excitatory connections while inhibitory neuronal populations have outgoing inhibitory connections. The connections marked with a “+/-” sign represent the connections that undergo synaptic plasticity: the connections between populations: “ES” to “EA”, “EA” to “EM-L”, and “EA” to “EM-R”. **Arrow “action”**: The activity within the “EM-L” and “EM-R” neuronal populations determines the action performed by the agent. Higher activity within either neuronal population will determine the agent to make a move to the left or to the right as exemplified by the large red arrows in the left box. **Green arrow “reward”**: for the Reinforcement Learning training strategy, the state of the environment is used to dictate the reward (or punishment) administered to the network for a correct (or incorrect) move.

### Simulations

We used the NEURON simulation environment (Carnevale and Hines 2006) with NetPyNE package (Salvador Dura-Bernal et al. 2019) for all modeling. NEURON simulates the firing of individual neurons based on the integration of input activation. Neurons are assembled into populations and into a connected network using the NetPyNE package that further coordinates the network simulation environment. The integration of the CartPole environment and the NetPyNE network was implemented in Python. The CartPole environment and the network simulation (the agent) is synchronized every time step T (50ms) in the following way:

- at the beginning of the time step, the environment is translated into neural activity in the input population (ES);
- for the duration of the time T, neurons are spiking based on induced activity;
- at the end of the time step (after 50ms), higher relative activation in the motor populations (EM-L, EM-R) determines the agent’s action.

In the following sections, we will present the setup of the pre-plasticity simulation: the excitatory/inhibitory neurons, the network inputs, the movement generation, and the weights initialization.

### Constructing a spiking neuronal network model to play CartPole

Our SNN model was adapted from one of our recent models (Anwar et al. 2021). To allow the SNN model to capture the game-state space reliably, we included 80 neurons in the sensory area (ES) with four subpopulations (20 neurons each), each to independently encode position, velocity, angle, and angular velocity (**Figure 1 right box, “ES”**). Each neuron was assigned to encode a different receptive field. Since the game’s goal was to balance the pole, that would require more precision in encoding sensory information near balanced states, around the absolute value of 0. To capture higher sensory precision utilizing smaller receptive fields around the balanced state and less precision utilizing larger receptive fields at peripheries, we assigned receptive fields to each neuron based on percentiles of a Gaussian distribution with a peak value of 0 centered around the 11th neuron. As such, lower indices neurons (neurons 1-10) encode negative values of the state-variables in decreasing order. Similarly, higher indices neurons (neurons 11-20) encode positive values of the state-variables in increasing order. All four ES populations were assigned receptive fields using Gaussian distributions with each input state’s expected mean and variance.

At each game-step, 4 ES neurons, one from each subpopulation, were activated, informing the SNN model about the game-state. To allow association between individual state-variable representing a game-state, we included 40 neurons in the association area “EA” (**Figure 1 right box, “EA”**), which received inputs from the ES neuronal population. Each neuron in EA was connected to motor areas EM-L and EM-R generating Right- and Left-actions (**Figure 1 “action” arrow**) by comparing the number of spikes in those populations (winner-takes-all). If both subpopulations have the same number of spikes, then a random move is performed.

To prevent hyperexcitability and depolarization-block (Anwar et al. 2021; Chadderdon et al. 2012; Neymotin et al. 2013), we included inhibitory neuronal populations (10 IA, 10 IAL, 10 IM, and 10 IML) and synapse weight normalization steps (detailed below).

### Integrate-and-Fire neuron

Individual neurons were modeled as event-driven, rule-based dynamical units with many of the key features found in real neurons, including adaptation, bursting, depolarization blockade, and voltage-sensitive NMDA conductance (Lytton et al. 2008; Lytton and Stewart 2006; Neymotin et al. 2011; Lytton and Omurtag 2007). Event-driven processing provides a faster alternative to network integration: a presynaptic spike is an event that arrives after a delay at a postsynaptic neuron; this arrival is then a subsequent event that triggers further processing in the postsynaptic neurons. Neurons were parameterized (**Table 1**) as excitatory (E), fast-spiking inhibitory (I), and low threshold activated inhibitory (IL). Each neuron had a membrane voltage state variable (V_m_), with a baseline value determined by a resting membrane potential parameter (V_rest_). After synaptic input events, if V_m_ crossed the spiking threshold (V_thresh_), the cell would fire an action potential and enter an absolute refractory period, lasting τ_AR_ ms. After an action potential, an after-hyperpolarization voltage state variable (V_AHP_) was increased by a fixed amount ΔV_AHP_, and then V_AHP_ was subtracted from V_m_. Then V_AHP_ decayed exponentially (with the time constant τ_AHP_) to 0. To simulate depolarization blockade, a neuron could not fire if V_m_ surpassed the blockade voltage (V_block_). Relative refractory period was simulated after an action potential by increasing the firing threshold V_thresh_ by W_RR_(V_block_-V_thresh_), where W_RR_ was a unitless weight parameter. V_thresh_ then decayed exponentially to its baseline value with a time constant τ_RR_.

**Table 1:** Parameters of the neuron model for each type.

| Cell type | $V_{rest}$<br>(mV) | $V_{thresh}$<br>(mV) | $V_{block}$<br>(mV) | $\tau_{AR}$<br>(ms) | $W_{RR}$ | $\tau_{RR}$<br>(ms) | $\Delta V_{AHP}$<br>(mV) | $\tau_{AHP}$ (ms) |
| --- | --- | --- | --- | --- | --- | --- | --- | --- |
| Excitatory<br>(E) | -65 | -40 | -25 | 5 | 0.75 | 8 | 1 | 400 |
| Inhibitory<br>(I) | -63 | -40 | -10 | 2.5 | 0.25 | 1.5 | 0.5 | 50 |
| Low-thresh<br>old<br>Inhibitory<br>(IL) | -65 | -47 | -10 | 2.5 | 0.25 | 1.5 | 0.5 | 50 |

$V_{rest}$ =resting membrane potential; $V_{thresh}$ =spiking threshold, $V_{block}$ =depolarization blockade voltage, $\tau_{AR}$ =absolute refractory time constant, $W_{RR}$ =relative refractory weight, $\tau_{RR}$ =relative refractory time constant, $\Delta V_{AHP}$ =after-hyperpolarization increment in voltage, $\tau_{AHP}$ =after-hyperpolarization time constant.

### Synaptic mechanisms

In addition to the intrinsic membrane voltage state variable, each cell had four additional voltage state variables V_syn_, corresponding to the synaptic inputs. These represent AMPA (AM2), NMDA (NM2), and somatic and dendritic GABAA (GA and GA2) synapses. At the time of input events, synaptic weights were updated by step-wise changes in V_syn_, which were then added to the cell’s overall membrane voltage V_m_. To allow for dependence on V_m_, synaptic inputs changed V_syn_ by dV=W_syn_(1-V_m_/E_syn_), where W_syn_ is the synaptic weight, and E_syn_ is the reversal potential relative to V_rest_. The following values were used for the reversal potential E_syn_: AMPA, 0 mV; NMDA, 0 mV; and GABAA, –80 mV. After synaptic input events, the synapse voltages V_syn_ decayed exponentially toward 0 with time constants τ_syn_. The following values were used for τ_syn_: AMPA, 20 ms; NMDA, 300 ms; somatic GABAA, 10 ms; and dendritic GABAA, 20 ms. The delays between inputs to dendritic synapses (dendritic GABAA) and their effects on somatic voltage were selected from a uniform distribution ranging between 3–12 ms, while the delays for somatic synapses (AMPA, NMDA, somatic GABAA) were selected from a uniform distribution ranging from 1.8–2.2 ms. Synaptic weights were fixed between a given set of populations except for those involved in learning (see **the “+/-” sign in Fig. 1 right box** and plasticity “on” in **Table 2**).

**Table 2:**
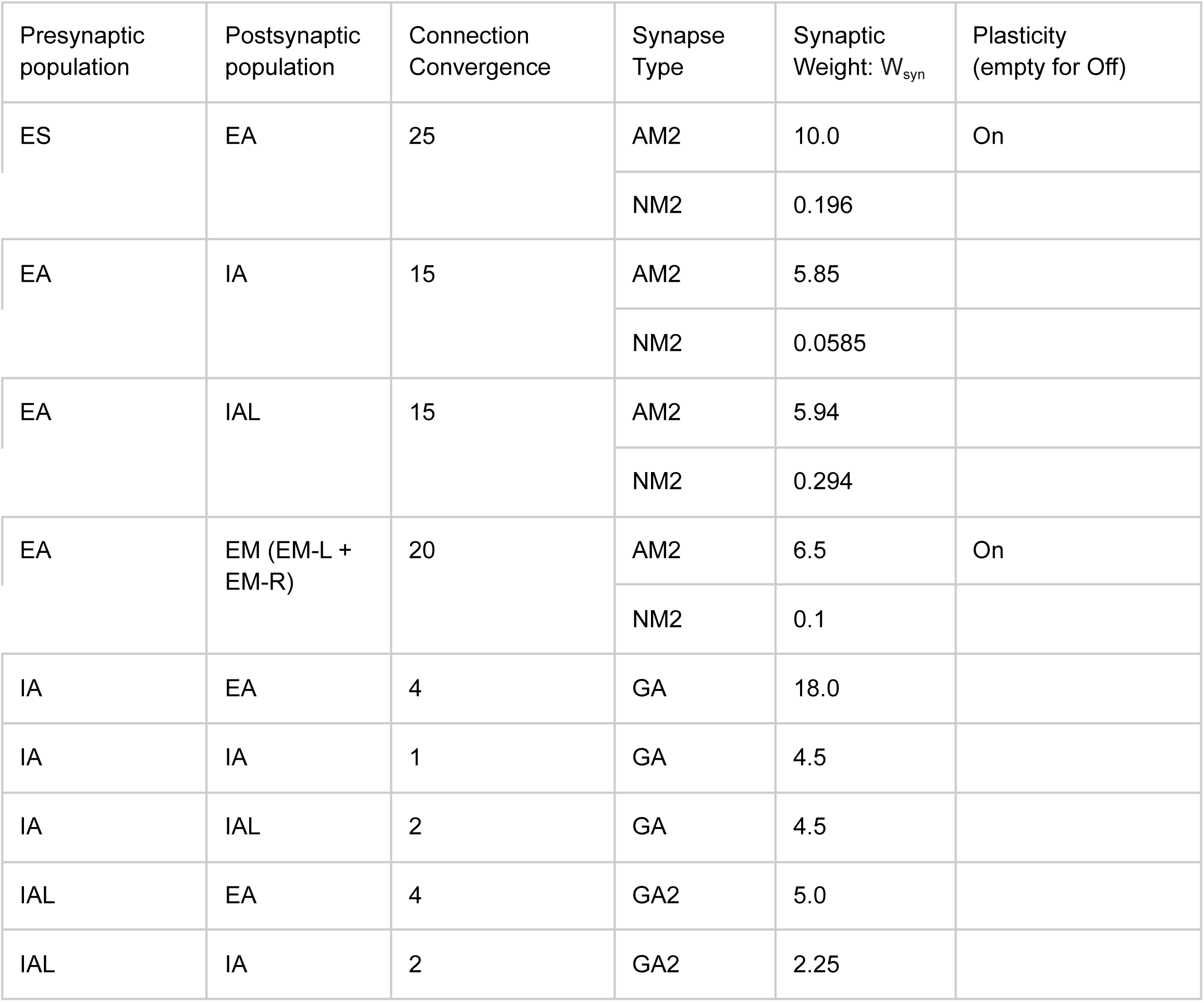
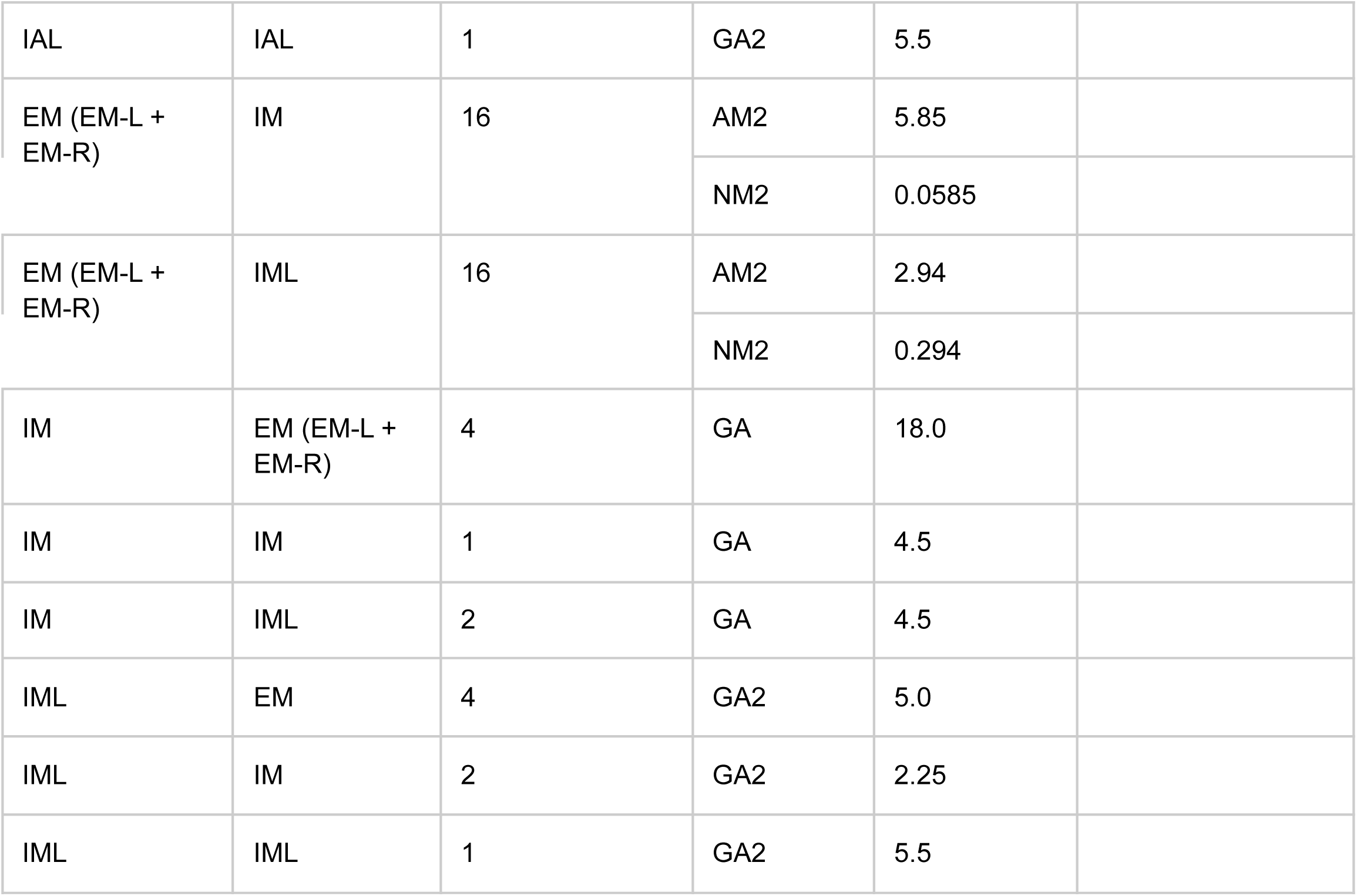
Initial connection weights.

| Presynaptic population | Postsynaptic population | Connection Convergence | Synapse Type | Synaptic Weight: $W_{syn}$ | Plasticity (empty for Off) |
| --- | --- | --- | --- | --- | --- |
| ES | EA | 25 | AM2 | 10.0 | On |
|  |  |  | NM2 | 0.196 |  |
| EA | IA | 15 | AM2 | 5.85 |  |
|  |  |  | NM2 | 0.0585 |  |
| EA | IAL | 15 | AM2 | 5.94 |  |
|  |  |  | NM2 | 0.294 |  |
| EA | EM (EM-L + EM-R) | 20 | AM2 | 6.5 | On |
|  |  |  | NM2 | 0.1 |  |
| IA | EA | 4 | GA | 18.0 |  |
| IA | IA | 1 | GA | 4.5 |  |
| IA | IAL | 2 | GA | 4.5 |  |
| IAL | EA | 4 | GA2 | 5.0 |  |
| IAL | IA | 2 | GA2 | 2.25 |  |
| IAL | IAL | 1 | GA2 | 5.5 |  |
| EM (EM-L + EM-R) | IM | 16 | AM2 | 5.85 |  |
|  |  |  | NM2 | 0.0585 |  |
| EM (EM-L + EM-R) | IML | 16 | AM2 | 2.94 |  |
|  |  |  | NM2 | 0.294 |  |
| IM | EM (EM-L + EM-R) | 4 | GA | 18.0 |  |
| IM | IM | 1 | GA | 4.5 |  |
| IM | IML | 2 | GA | 4.5 |  |
| IML | EM | 4 | GA2 | 5.0 |  |
| IML | IM | 2 | GA2 | 2.25 |  |
| IML | IML | 1 | GA2 | 5.5 |  |

### The Neuronal Weights

The neurons are organized into three overall layers: Sensory, Association, and Motor (**Figure 1 right box**). The sensory layer consists of the excitatory sensory neuronal population (ES) which contains a total of 80 excitatory neurons. The association layer consists of 40 excitatory neurons in the excitatory association neuronal population (EA), 10 fast-spiking inhibitory neurons in the inhibitory association neuronal population (IA), and 10 low threshold activated inhibitory neurons in the “low” inhibitory association neuronal population (IAL). Similarly, the motor layer consists of 40 excitatory neurons in the excitatory motor neuronal population (EM), 10 fast-spiking inhibitory neurons in the inhibitory motor neuronal population (IM), and 10 low threshold activated inhibitory neurons in the “low” inhibitory association neuronal population (IML). Furthermore, the EM neuronal population is split into 20 neurons associated with left movements (EM-L) and 20 neurons associated with right movements (EM-R).

The weights between populations were adjusted to allow reliable transmission of spiking activity across different layers/areas of the SNN model. Each row in **Table 2** describes the synaptic connectivity between two different neuronal populations (no self connectivity) and follows the same diagram of neuronal connectivity described in **Figure 1**. Neurons belonging to the presynaptic population have axons that project to neurons in the postsynaptic population. Each neuron projects to a fixed number of post-synaptic neurons, a constant defined as the connection convergence (**Table 2**). The individual neuronal connections are picked randomly at the initialization process and different random seed values generate different connections, hence different networks. The excitatory neuron have both AMPA (AM2) and NMDA (NM2) synaptic connections, while the inhibitory neurons either have somatic GABAA synapses (GA) for fast-spiking inhibitory neurons or have the dendritic GABAA synapses (GA2) for low threshold activated inhibitory neurons. The synaptic weight W_syn_ for each neuronal connection was picked based on previous studies and fine-tuned on this network to start with a biologically reasonable spiking pattern (2-20 Hz).

Some of the synapse weights can be changed throughout the course of the training simulation as they undergo synaptic plasticity. For this work, we limited synaptic plasticity to AMPA synapses between excitatory populations (**Table 2: Plasticity column**). We found that this limitation is not hindering the network’s ability to learn the dynamical behavior of the CartPole problem, and it rather simplifies the network analysis. To have a consistent examination between different training strategies, we used the same initialization methods and plastic synapses as defined in above and in **Table 2**.

### Training strategies

#### Spike-timing dependent Reinforcement Learning (STDP-RL)

To train the neuronal networks, we used an existing STDP-RL mechanism, developed based on the distal reward learning paradigm proposed by Izhikevich (Izhikevich 2007), with variations used in spiking neuronal network models (Neymotin et al. 2013; Chadderdon et al. 2012; Salvador Dura-Bernal et al. 2016; Chadderdon and Sporns 2006; Anwar et al. 2021). Our version of STDP-RL (**Figure 2A**) uses a spike-time-dependent plasticity mechanism together with a reward or punishment signal for potentiation or depression of the targeted synapses. When a postsynaptic spike occurred within a few milliseconds of the presynaptic spike, the synaptic connection between this pair of neurons became eligible for STDP-RL and was tagged with an exponentially decaying eligibility trace. An exponentially decaying eligibility trace was included to assign temporally distal credits to the relevant synaptic connections. Later, when a reward or a punishment was delivered before the eligibility trace decayed to zero, the weight of the tagged synaptic connection was increased or decreased, depending on the ‘critic’ value and sign, i.e., increase for reward or decrease for punishment. Since we used an eligibility trace with an exponent of 1 in this work, the change in synaptic strength was constant at the time of the critic’s delivery.

**Figure 2:**
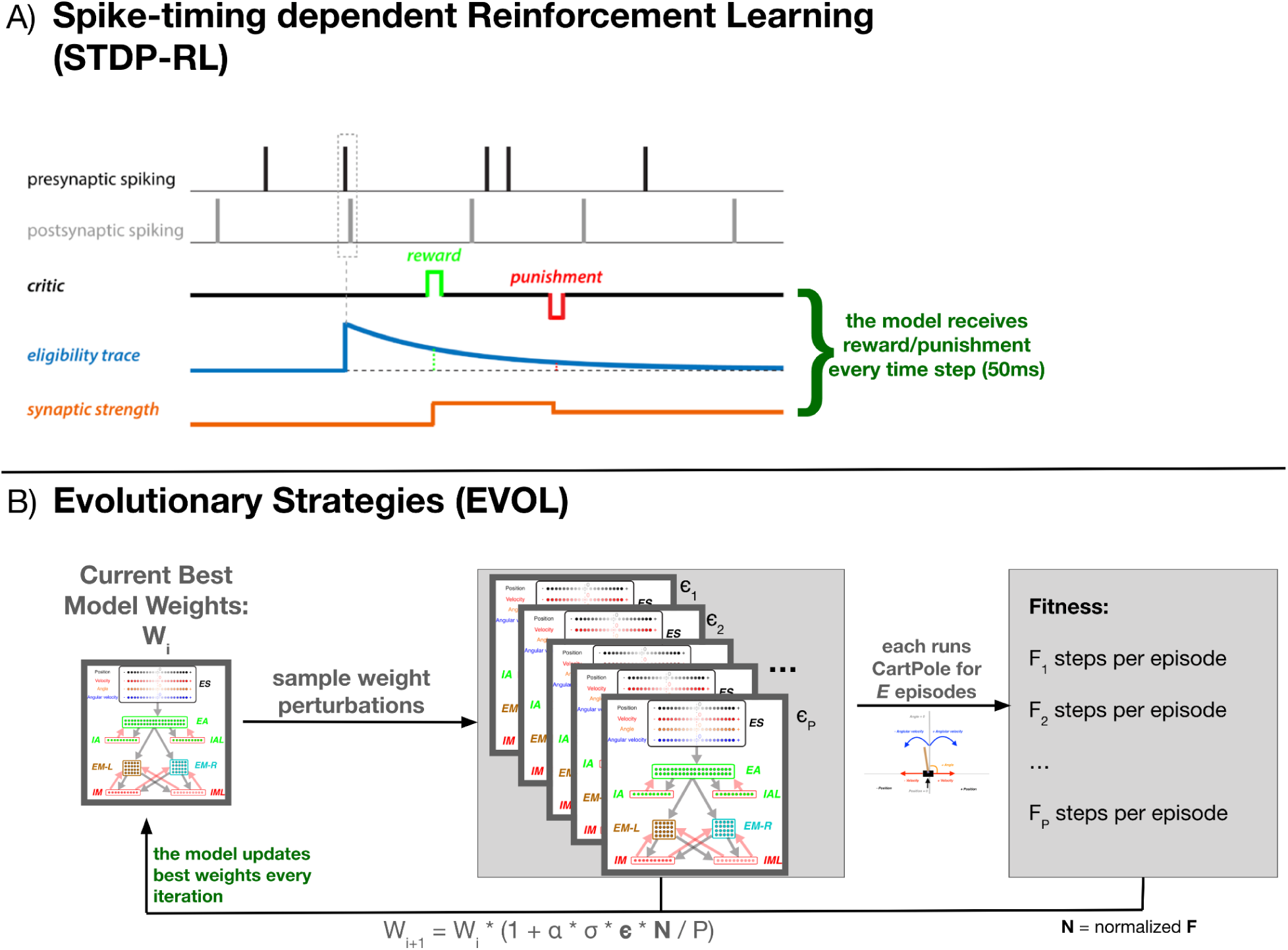
Training SNN using STDP-RL and EVOL strategies. **A)** In STDP-RL, when a postsynaptic neuron produced a spike within a short-time interval of a presynaptic spike, the synapse between the pair of neurons was tagged with a decaying eligibility trace. The tagged synapse was strengthened or weakened proportional to the value of eligibility trace for a reward or punishment, respectively. **B)** Schematic showing the steps of evolutionary strategies training algorithm (EVOL).

Traditionally, when using STDP-RL for learning behavior, all plastic synaptic connections in the neuronal network model are treated equally (*non-targeted STDP-RL*). This strategy considers that the underlying causality between pre and postsynaptic neurons and the associated reinforcement automatically changes only relevant synaptic connections. On top of the traditional STDP-RL approach, we used two recently developed versions of targeted reinforcement by selectively delivering reward and punishment to different subpopulations of the Motor population (EM) (Anwar et al. 2021). In the first variation (*targeted RL main*), we delivered reward or punishment only to the neuronal subpopulation that generated the action (EM-LEFT or EM-RIGHT). In the second variation (*targeted RL both*), we additionally delivered opposite and attenuated reinforcement to the opposite-action neuronal subpopulation. Both targeted methods ensured that the learning was specific to the part of the circuit which generated the action. Moreover, we explored delivering (attenuated) ‘*critic*’ values to the neuronal populations one synapse away from those directly generating motor actions (EA population). We used and evaluated all six STDP-RL mechanisms (three targeted RL versions X two nonMotor RL versions) for learning performance during hyperparameter search (see below for details). In all cases, although there is evidence of plasticity involving inhibitory interneurons in vivo(Anwar et al. 2017; Vogels et al. 2011), for the sake of simplicity STDP-RL was only applied between excitatory neurons.

#### Critic

For STDP-RL, the model relies on a critic to provide essential feedback to the model’s actions (**Figure 2A**). For CartPole, we picked a critic that responds positively to movements that bring the vertical pole closer to a balanced position. We computed a loss for each position, determined by the absolute values of the angle and the angular velocity input states. The critic’s returned value will be the difference between the loss of the previous state and the loss of the current state. If the loss between the following states increases due to the agent’s move, then the critic will return a negative value, corresponding to a punishment. Similarly, a decrease in loss will return a positive critic value, corresponding to a reward. If the agent could not decide on a motor move due to identical spiking activity in both subpopulations, then a constant negative punishment is returned. Additionally, as the critic is dominated by punishment at the beginning of training, to avoid the weights decreasing to zero, the model needs an associated boost in positive rewards (η_*positivity*_). The critic is finally capped at a minimum and maximum value to keep rewards within the interval: [− *max_reward, max_reward*].

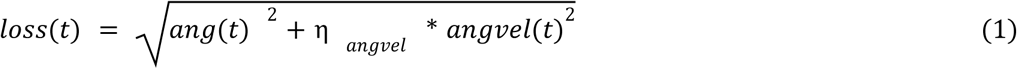

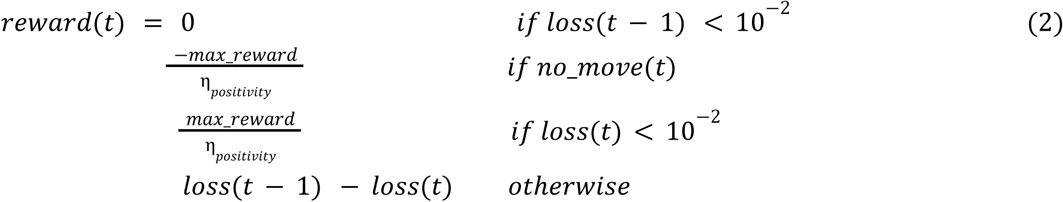

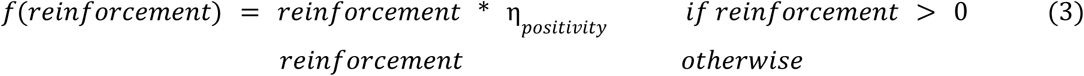

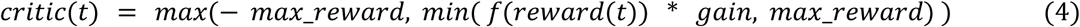

Where:

- *ang*(*t*)and *angvel*(*t*)represent the input states at time step t;
- η_*angvel*_ represents the angular velocity bias, used to balance the dominance of each input state;
- η_*positivity*_ represents the positivity bias to reinforce rewarding behavior;
- *max_reward* is the maximum reward used, fixed at 1. 0 throughout the whole experiment;
- *gain*represents a final multiplier that increases the absolute reward value.

The critic was implemented as a crude reinforcer of synaptic plasticity, working in conjunction with STDP events. As we found in the HyperParameter search (described below), most of the hyperparameters of the critic have little influence over the final performance of the model, and we believe that many different critic functions would have been suitable for our analysis. More importantly, for synaptic weight normalization, the critic values are further modulated by output gain and homeostatic gain control, as described below.

#### Hyperparameter Search

We first trained our SNN model using the STDP-RL parameters’ values as used in earlier studies (S. Dura-Bernal et al. 2017) and found that the model did not perform very well since it could not learn to balance the CartPole for 50 steps per episode (averaged over 100 episodes). The low performance indicated that these parameters might not be optimal for training with STDP-RL. To find an optimal training strategy, we perturbed parameters of our STDP-RL training setup. We identified nine hyperparameters to optimize(**Supplementary Table 1**), related specifically to the STDP-RL mechanism that are somewhat independent of each other. Out of the nine: three parameters influence the timing and weight update of the STPD-modulated AMPA synapse(AMPA-RLwindhebb, AMPA-RLlenhebb, AMPA-RLhebbwt); four parameters determine the area of effect of STDP (Targeted_RL_Type, Non_Motor_RL, Targeted_RL_Opp_EM, Targeted_RL_Non_Motor); and two parameters influence the reinforcement derived from the critic(Critic Positivity Bias η_*positivity*_, Critic Angv Bias η_*angvel*_). To identify the best combination of the hyperparameters for training the network, we ran multiple random hyperparameter searches on those nine hyperparameters (**Supplementary Table 1**).

For the first hyperparameter search, we evaluated the nine parameters above by training networks with random choices of those hyperparameters. From the 7200 possible combinations, we sampled 50 combinations and trained randomly-initialized models using STDP-RL for 500 seconds in simulation time. We evaluated the performance of those models based on the average steps per episode over 100 episodes during training(**Supplementary Figure 1**). Only the Targeted_RL_Type hyperparameter performance distributions showed a significant deviation from the mean performance (ANOVA p-value < 10^−5^), but it failed the homogeneity of variance assumption test.

For the second step of the hyperparameter search, we continued training from the best four models from the previous step, and continued evaluating the nine parameters. For this step, we had 6912 combinations from which we sampled 54 combinations that we trained for 2000 seconds in simulation time. Similarly, we evaluated the performance of those models based on the average steps per episode over 100 episodes during training(**Supplementary Figure 2**). The initialization model choice displayed a significant contribution to the final model performance (ANOVA p-value = 0.00012). Moreover, the hyperparameter defining the maximum time between pre- and postsynaptic spike (AMPA-RLwindhebb), also showed a minimal deviation from the mean performance (ANOVA p-value = 0.027).

For the third step of the hyperparameter search, we continued training from the best four models from the previous step, and continued evaluating the nine parameters. For this step, we had 1728 combinations from which we sampled 56 combinations that we trained for 2000 seconds in simulation time. Similarly, we evaluated the performance of those models based on the average steps per episode over 100 episodes during training(**Supplementary Figure 3**). The initialization model choice displayed a significant contribution to the final model performance (ANOVA p-value = 0.00023). Moreover, the hyperparameter defining the choice of activating plasticity for the synapses within the *ES* → *EA* pathway (Non_Motor_RL), showed a minimal deviation from the mean performance (ANOVA p-value = 0.005). This latter finding was further tested in detail by training multiple models with and without *ES* → *EA* plasticity (see Results).

Most of the hyperparameters had a minimal effect on the final training performance for the STDP-RL model as there is no significant difference between the performance of models with different hyperparameter values. Interestingly, the hyperparameter search revealed better preference when using the “*targeted RL both”* paradigm. These findings suggest that targeted plasticity of specific motor areas could enhance the learning ability of the model, consistent with earlier findings((Anwar et al. 2021; Patel et al. 2019; Hazan et al. 2018; Chadderdon et al. 2012)).

#### Training Protocol for the STDP-RL model

Since we couldn’t establish a best choice for each of the hyperparameters, we used the training protocol of the best model resulting from the third hyperparameter step. Hence, we trained our STDP-RL models with a sequence of the hyperparameter values for different durations (500s, 2000s, 2000s). Continuing training after the 4500s mark, we used the hyperparameters of the third configuration. The exact hyperparameter values used are highlighted in **bold** in **Supplementary Table 1**.

#### Evolutionary Strategies

The Evolutionary Strategies (EVOL) algorithm has been shown to be an effective gradient-free method for training ANNs in reinforcement learning control problems (Salimans et al. 2017). Here we adapt this learning technique to SNNs to solve the CartPole problem by procedurally adapting the plastic weights of the SNN. Our method progressively adapts only the weights and not the delays as it was used in previous Evolutionary Strategies for SNNs (Altamirano et al. 2015). It should be noted that in the ANN implementation of EVOL weights are allowed to be unrestricted in value, so additive weights were used. As SNNs don’t have negative weights, we instead use a multiplicative noise, i.e. we increase or decrease the weights by a randomly selected percentage. In this way we are able to restrict the SNN weights to valid positive values while still effectively searching all possible parameterizations.

Formally our EVOL algorithm (**Figure 2B**) consists of the following steps to change the weight for each synapse, performed in parallel for the whole model: (1) at iteration i, keep track of current best synapse weight: w_i_; (2) sample a population(P) of weight perturbations ϵ_1..P_ from the normal distribution; (3) for each weight perturbation (ϵ_j_), evaluate the whole network weights (w_i_ * (1 + σ * ϵ_j_)) on the CartPole environment for a fixed number of episodes (X); (4) measure the fitness (F_j_) as the average count of steps achieved during the X episodes; (5) Normalize population fitness values (N_j_) by subtracting the population mean fitness and dividing by the population mean standard deviation (6) modulate the weight perturbations based on the normalized fitness and derive a new best synapse weight:

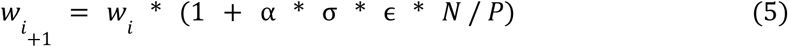

Where α is the learning rate, σ is the noise variance, and **ϵ** and **N** are the vector representations of the weight perturbations for each synapse and the normalized fitness, respectively. We only update the weights that undergo synaptic plasticity (**Table 2**). The weights are initialized (w_0_) as the same initial weights we used for the STDP-RL model. In this case, STDP was fully deactivated and the EVOL training procedure updated the synaptic weights every iteration.

We trained using the EVOL algorithm on multiple models for 1500 iterations and a population of P=10 with synapse weight perturbations of σ=0.1 variance. We used a learning rate of α=1.0. We used 5 and 10 episodes (value X above) during fitness evaluation. The model weights were initialized based on the best and worst models trained with STDP-RL.

#### Synaptic weight normalization

Training the model with STDP-RL, the synaptic weights tend to increase without bound, leading to epileptic activity (M. S. Rowan, Neymotin, and Lytton 2014). To avoid this behavior, we incorporated biologically-realistic normalization methods (Sanda, Skorheim, and Bazhenov 2017; M. Rowan and Neymotin 2013). Apart from the inhibition mechanisms described earlier, we used the following techniques: balancing of synaptic input, output balancing, homeostatic gain control.

To balance the combined synaptic input to each neuron, the total reception weight (defined as the sum of synaptic weights onto each postsynaptic neuron from multiple presynaptic neurons) is normalized to initial values every 25 time steps. As this procedure keeps neuronal inputs constant, it either decreases the weights of specific unused synapses or promotes beneficial synapses, creating synaptic competition.

To prevent synapses from overwhelming the network with ever-increasing rewards, each synapse’s reinforcement is scaled by the change in the neuron’s total transmission weight (defined as the sum of synaptic weights from a presynaptic neuron onto multiple postsynaptic neurons). Hence, a synapse with a high transmission weight compared to initialization will scale down the reward value and scale up the punishment value during STDP-RL events. Conversely, a synapse with a low transmission weight compared to the initialization will increase STDP-RL reward values while decreasing STDP-RL punishment values. For this normalization step, we cap the maximum scaling factor for promoting rewards or punishments to 0.1 for scaling down and 2.0 for scaling up.

Homeostatic gain control is a normalization method of keeping neuronal populations within a target spiking rate(measured over 500 time steps). The method updates the target population synaptic inputs every 75 timesteps after checking if the firing rate is different from the target firing rate (5.5 Hz for EA and 6.0 Hz for EM). The homeostatic gain control normalization method doesn’t update the synaptic weights directly but rather scales the expected total reception weight (by 0.01%) that is later used for normalization during the balancing of synaptic inputs (see above). This procedure has the effect of reducing high neuronal activity and promoting baseline activity (Sanda, Skorheim, and Bazhenov 2017; M. Rowan and Neymotin 2013)). As this update is infrequent (every 75 time steps) and has a small indirect update (scaling by 0.01%), over a large amount of training, this method has an effect similar to the network noise.

#### Validation, Testing, and All-Inputs datasets

For testing the trained models, we used two separate datasets, the validation dataset for selecting the best model timepoint and the testing dataset for reporting the final model performance. For those two datasets, we set a seed value for the OpenAI gym environment in order to fix the episodes to consistently get the same episode initialization. This environment seed(S_r1_, S_r2_, …, S_r9_) should not be confused with the model seed(Seed-6, Seed-3, …) that we use to randomly initialize the neuronal connections. The episodes are fixed for our datasets when they have the same initial starting conditions for all four environment parameters (position, velocity, angle, angular velocity). While training the models, the episodes are not fixed and are randomly selected for each new training instantiation. To select the best saved time-point of a trained model, we evaluate the model with weights at different time-points throughout training using a validation dataset with 100 fixed episodes. Once the best time-point of a model is picked, we evaluate and report the performance on the testing dataset which contains another 100 fixed episodes. For analysis, we selected some of the fixed episodes for further evaluation.

As each input parameter space is discretized into activation of twenty individual neurons, the total possible combinations of inputs is 20^4^. We tested the models on all the possible combinations, the *All-Inputs dataset*, by activating each possible input combination at a time. Sensory activations were interleaved with blank periods (no stimulation) to make sure that the effect of activation does not bleed into next sensory activation.

#### Software

All modeling and analysis source code is available on github at the following repository location https://github.com/NathanKlineInstitute/netpyne-STDP.

## Results

Like in any multilayer artificial neural network, the biggest challenge of training SNN models to perform sensory-motor behaviors is to come up with a network design and to choose the right set of learning parameters resulting in optimal performance. This leaves model developers with many possibilities. Instead of trying many network configurations, here we constrained our network to a minimal configuration (see Methods) based on our previous study (Anwar et al. 2021) where the SNN models learned to play a racket-ball game showing robust performance using several STDP-RL setups. Our model included 3 layers of excitatory neuronal populations: *ES* representing a sensory area, *EA* representing an association area, and *EM* representing a motor area with probabilistic inter-area connections and fixed initial synaptic synaptic weights (see Methods, “The Neuronal Weights”). Inhibitory neurons in each area were included to prevent network hyperexcitability after increased synaptic weights from learning. The activity in the network propagated in a single direction: *ES* → *EA* → *EM*. To compare different learning strategies, we further constrained our network to two sets of plasticity based configurations: 1. only *EA* → *EM* synapses are plastic and 2. Both *ES* → *EA* and *EA* → *EM* synapses are plastic. Using these two plasticity constraints, we trained our SNNs with STDP-RL and EVOL methods, evaluated their performances and dissected the circuits post-training to compare the underlying dynamics resulting in diverse behavioral performances.

### Training Multilayered SNN models using STDP-RL improved performance

We first trained our SNN model using STDP-RL while allowing only *EA* → *EM* synaptic connections to adapt. Since the number of neurons was small in each layer and inter-layer connections were probabilistic, there was a possibility that each initialization would result in a slightly different connectivity pattern, which might affect the learning capabilities of the model. To assess the effect of network initializations on the training, we trained the model using 10 different initializations (random number generator seeds). Each training session included a large number of CartPole episodes where each episode was randomly initialized. Such random initializations of the game exposed the model to different sensory states representing the game environment and required the model to learn different action strategies during each episode. Learning sensory-action strategy for an episode resulting in a higher performance does not necessarily mean that the model will perform equally well during the next episode, as evident by large fluctuations in the performance during the training (blue dots in **Figure 3A**). Such variability in the performance makes it difficult to accurately quantify the learning performance of the model. Therefore, we averaged the performance over 100 episodes to quantify the learning quality of the model (**Figure 3A**). We continued training each model until the performance converged or started decreasing, which typically required training over simulation durations of 25000 seconds. We observed a strong effect of the model initialization on its performance, as the maximum performances of the models during training ranged from 75 to 157, with a mean of 118 steps per episode (**Figures 3B and E**).

**Figure 3:**
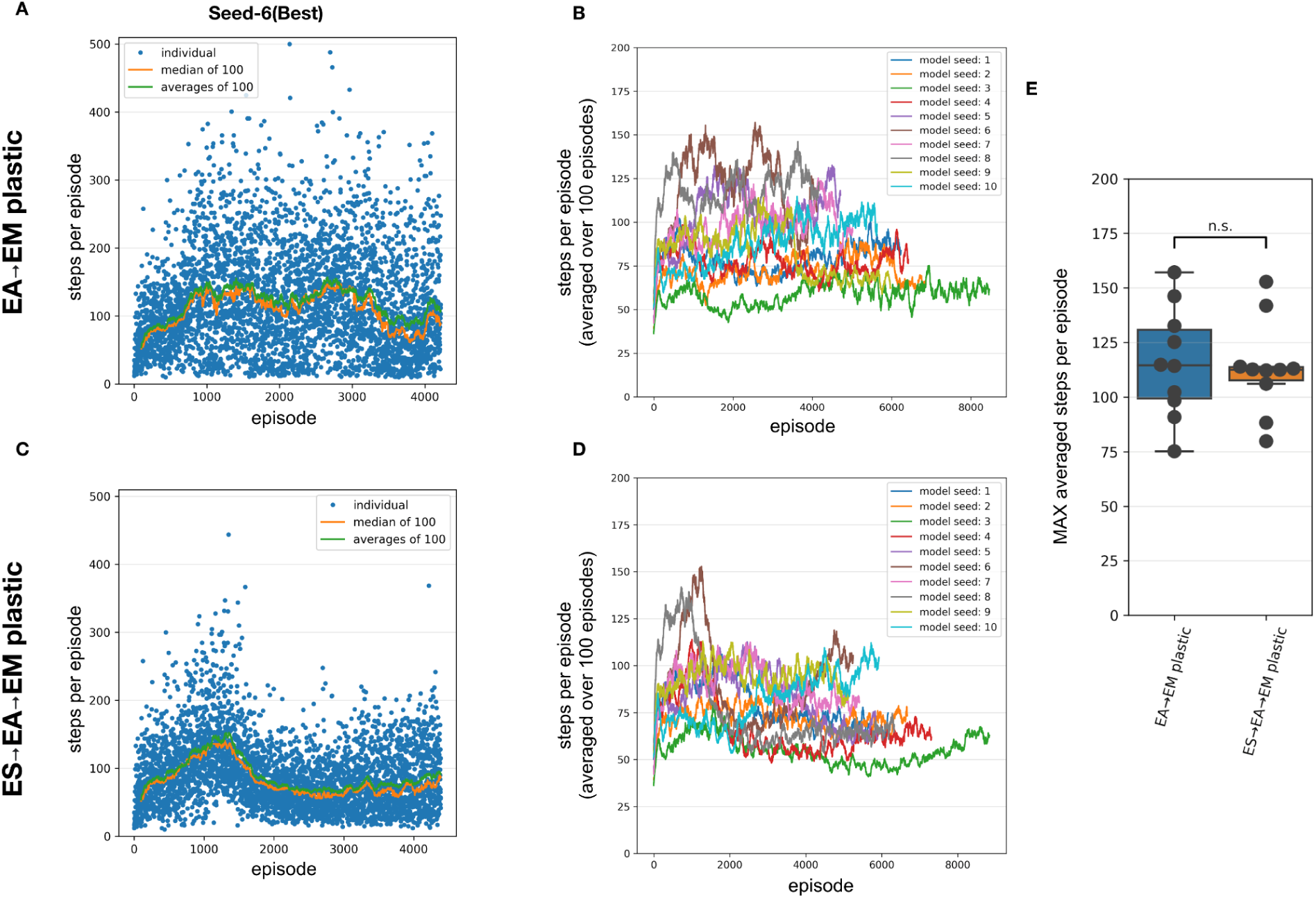
Performance of the SNN models when training with STDP-RL. **(A-B)** Training SNN model with only EA → EM plastic. **(A)** Training performance (defined as the length of time the model could keep the pole balanced) of a model initialized using Seed-6. Each blue dot shows performance during individual episodes. The green and orange lines represent the averages and the median performances respectively. **(B)** Training performance of the 10 randomly initialized models. All models trained for 25000 sec of simulated time. Out of the tested models, the seed-3 model was the worst performer(green line), while the seed-6 model was the best performer(brown line and in **A**). **(C and D)** Same as **A** and **B** respectively, with both ES → EA and EA → EM synaptic connections plastic. **(E)** Comparing the peak average performance of 10 randomly initialized models with only EA → EM plastic synapses vs both ES → EA and EA → EM plastic synapses. The box plot represents the distribution of maximum averages over 100 steps of the models. Each dot of the scatter plot represents the maximum average over 100 steps of one of the randomly initialized models. Comparing the two distributions using a 1-way ANOVA, we obtained a non-significant result (p=0.8), showing that the two types of model plasticities have similar performance.

To further investigate whether allowing synaptic connections between the sensory and the association layer (*ES* → *EA*) to learn via STDP-RL in addition to *EA* → *EM* synaptic connections will enhance the capability of our model to perform better, we repeated training simulations using 10 different network initializations (same random number generating seeds as in **Figure 3B**). The models showed large variance in the performance from episode to episode during training as well as among themselves similar to the models with only *EA* → *EM* plasticity (**Figure 3C** vs **Figure 3A**). Although for many seeds, the peak performance was more consistent (**Figure 3E** right bar) regardless of varying performance trajectories (**Figure 3D**). Surprisingly, overall, the performance did not improve significantly (**Figures 3B, D and E**). Note that we used the same learning parameters in these simulations as in the models where only *EA* → *EM* synaptic connections were plastic (parameter values optimized using hyperparameter search, see Methods). Since these models did not perform better than the models with only *EA* → *EM* plasticity, we only considered models with plasticity in the *EA* → *EM* pathway for the remainder of the manuscript when comparing to EVOL based models, and analyses.

As shown in **Figure 3**, we observed a large variability in performance from episode to episode due to the dynamically evolving sensory-state of the environment across episodes. It remains unclear whether the performance will sustain after learning is stopped and weights are frozen. To test post-training performance, we further evaluated only two models with unique weights based on their performance during training; the best performing model (seed-6 in **Figure 3B**) and the worst performing model (seed-3 model, **Figure 3B**). We evaluated the selected models using 100 episodes with the same game initiatializations for both models, and compared them to the model before training and to a random choice null model (**Figure 4**). During training, both models improved their performance throughout STDP-RL training as we see an increase from 19 to 130 median steps per episode for the Seed-6(best) model, and from 19 to 53 median steps per episode for the Seed-3(worst) model. These performances were better than before training and compared to the random choice null models (**Figure 4**). Although the performance remained suboptimal (i.e. 500 steps per episode), both models showed enhanced performance after training (**Figure 4**). One-way ANOVA on the logarithm of the model performances before and after training resulted in p < 10^−10^.

**Figure 4:**
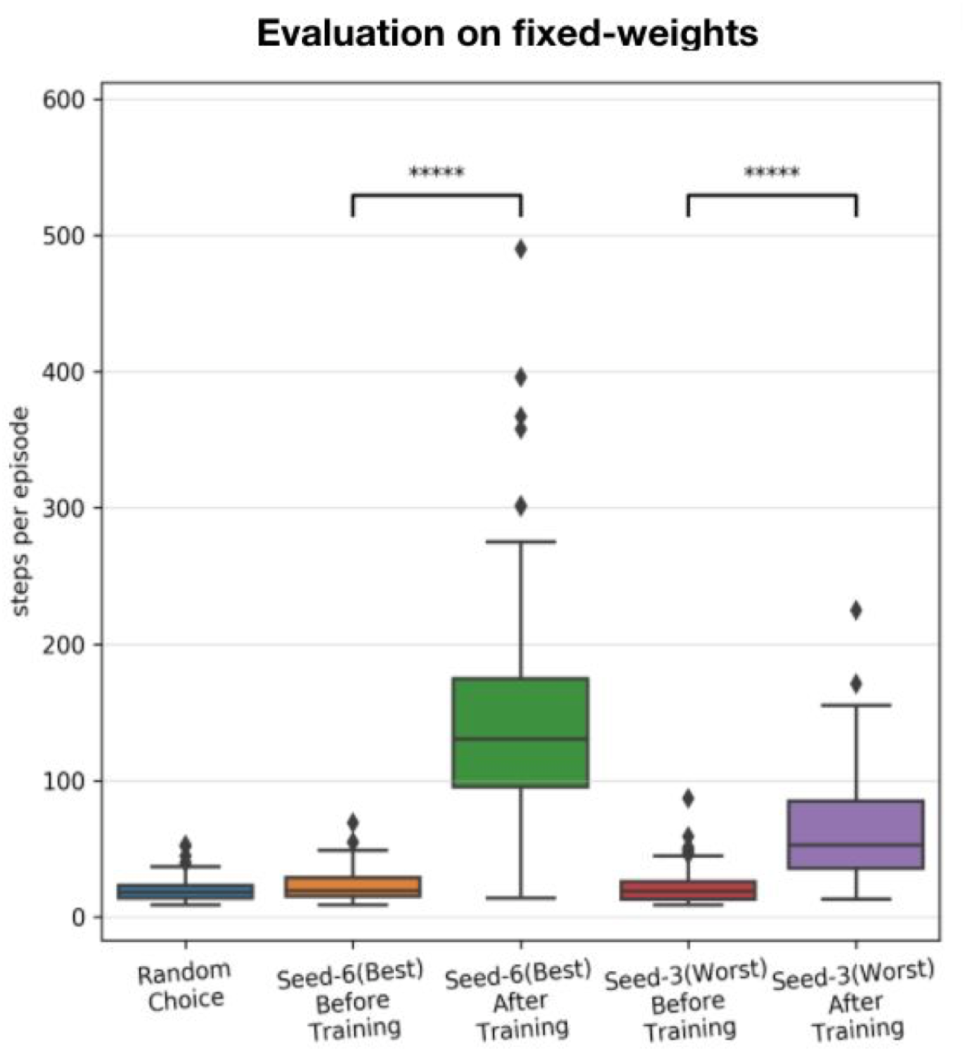
Performance of the SNN models after training with STDP-RL. The distribution of performance (steps per episode) calculated before and after training, on the testing dataset, using fixed weights of the seed-6 (best) and seed-3 (worst) models. The 5-star (*****) denotes p < 10^−10^ when comparing the two distributions with a one-way ANOVA.

### Multilayered SNN models trained using EVOL achieve optimal performance

Variability in performance due to different network configurations resulting from different network initializations suggests that the performance of these models is sensitive to the initial synaptic connections and corresponding weights. Although synaptic weights between two populations of neurons in neocortex may vary greatly, in modeling SNNs the initial weights are often drawn from a normal distribution around arbitrarily chosen and tested values, which may bias the ability of the network to learn a behavior. Evolutionary algorithms can allow parsing the weights of synaptic connections over a large range of values, where the subsequent performance evaluation of each set of synaptic weights can guide the model to evolve along complex and highly nonlinear trajectories, eventually producing models with better performance. To that end, we next evaluated the potential of our models to learn sensory-motor behavior utilizing the EVOL algorithm.

Similar to training using STDP-RL, we first trained our SNN model using the EVOL algorithm with plasticity limited to the *EA* → *EM* pathway. While the STDP-RL training performance plateaued before reaching an average of 160 steps per episode, some of the EVOL models performed better than 200 steps per episode (**Figures 5B and E**). We observed relatively smaller variance in performance from one iteration to the next one (**Figure 5A**). One reason for the low variance in performance across iterations could be due to the averaging performance of 50 episodes (5 episodes per perturbation) per iteration compared to showing performance of each individual episode in STDP-RL learning models (**Figure 3A**). The performance of our EVOL-trained models increased further when using SNN models with both *ES* → *EA* and *EA* → *EM* plastic synaptic connections. In fact, the average performance increased to optimal performance of 500 steps per episode (**Figures 5C, D and E**). We also observed further reduction in the variance of performance across iterations (**Figures 5C vs 5A**). Not only did EVOL produce optimal performance, but all the models rapidly learned the task and achieved high performance (>400 steps per episode) in roughly 250-500 iterations (**Figure 5D**). Note that during each iteration, EVOL generated *P (=10)* perturbations (each has a performance value) and running *E* (=5) episodes for each perturbation, for a total of *P * E (=50)* episodes. We display the performance minimum, average, and maximum for each iteration for the model with the Seed-3 random initialization (**Figures 5A and C**). Since the models with both *ES* → *EA* and *EA* → *EM* plastic synapses performed better than the models with only *EA* → *EM* plasticity, we only considered models with *ES* → *EA* and *EA* → *EM* plastic synapses in the remaining manuscript for the comparison to STDP-RL based models and the analysis. When we tested the selected models (trained using seed-6 and seed-3) using 100 episodes with the same game initiatializations, both models were able to hold the pole vertically up for an average of ∼500 steps per episode (**Table 3**).

**Table 3:** Performance comparison of training strategies on the testing dataset.

| Model | Average on Test Dataset | Median on Test Dataset | Trained Episodes | Iterations | Population |
| --- | --- | --- | --- | --- | --- |
| Seed-6 after initialization | 23.38 | 19.5 | 0 |  | - |
| Seed-6 after STDP-RL training | 144.67 | 130.5 | 4226 |  | - |
| Seed-6 after EVOL training | 499.42 | 500.0 | 80000 | 1600 | 10 |
| Seed-3 after initialization | 23.04 | 19.0 | 0 |  | - |
| Seed-3 after STDP-RL training | 64.14 | 53.0 | 8577 |  | - |
| Seed-3 after EVOL training | 499.09 | 500.0 | 80000 | 1600 | 10 |

**Figure 5:**
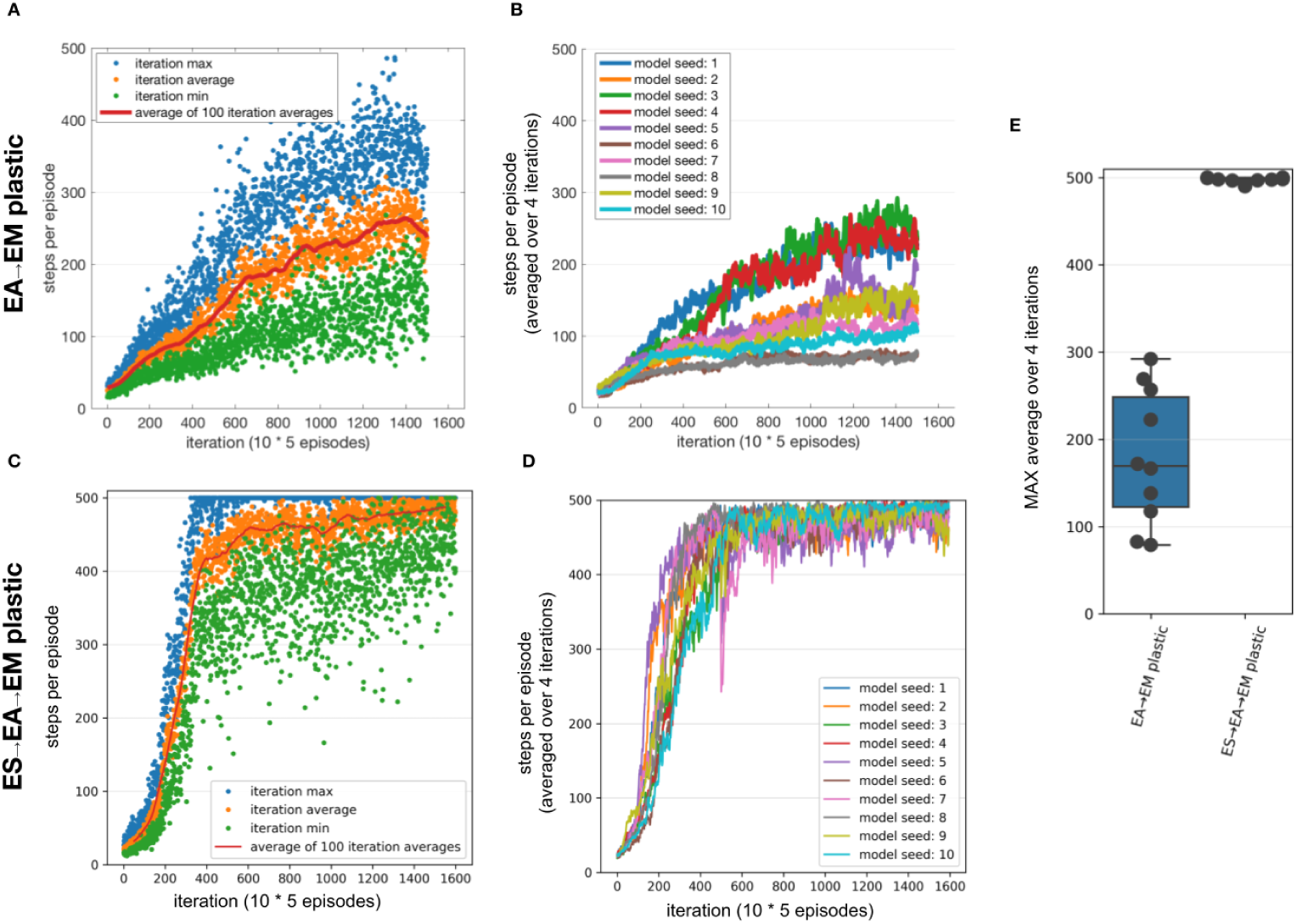
Performance of the SNN model when training with EVOL. **(A-B)** Training SNN model with only EA → EM plastic. (A) Training performance (defined as the length of time the model could keep the pole balanced) of a model initialized using Seed-3. During each iteration, 10 models with unique weights were evaluated on 5 episodes. Each red dot shows an average performance of (10 models x 5 episodes/model = 50 episodes), whereas each green dot and blue dot show minimum and maximum performance of 50 episodes. **(B)** Training performance of 10 randomly initialized models. All models trained for the same number of iterations as opposed to the same amount of simulated time. **(C and D)** Same as A and B respectively, with both ES → EA and EA → EM synaptic connections plastic. **(E)** Comparing the peak average performance of 10 randomly initialized models with only EA → EM plastic synapses vs both ES → EA and EA → EM plastic synapses. The box plot represents the distribution of maximum averages over 100 steps of the models. Each dot of the scatter plot represents the maximum average over 100 steps of one of the randomly initialized models.

Comparing with the STDP-RL training strategy, with which the model reached an average of 118 steps per episode over 3931 episodes on average, the EVOL models with*ES* → *EA* and *EA* → *EM* both plastic, used on average 145 iterations (for a total of 7250 training episodes) to reach the same performance level. While STDP-RL training was provided with frequent rewards from a hand-tuned critic, the EVOL training only used episodic fitness, a much sparser feedback signal. Although the STDP-RL models achieved an improved and sustained performance, we found EVOL strategy to be better suited for solving the CartPole problem.

### Variability in the performance of models trained using STDP-RL is partially related to the differences in the sensory environment (game initializations)

To demonstrate that the model learned the behavior and did not forget it, we earlier compared two different STDP-RL models (with different seeds) with the random-action generating and untrained models (**Figure 4**) and found variable yet sustained learning in the STDP-RL models. Negligible variance in the performance was also observed for models trained using EVOL. As demonstrated earlier (**Figures 3, 4 and 5; Table 3**), this performance variability could be related to the initialization of synaptic weights or the presence/absence of synaptic connections between specific pairs of neurons or on the initial game-state (note that we reset the game-states at the end of each episode). To test the latter possibility, we handpicked nine unique initial game-states based on the performance of each model on the initial evaluation. For each initial game-state, we repeatedly simulated each trained model over 25 episodes (**Figure 6**; note that at the end of each repeated episode, only the game-state was reset to the associated initial game-state, and the model neuron states were not reset). In **Figure 6**, Performance is shown only for STDP-RL models because EVOL models after training performed almost perfectly i.e ∼500 steps per episode. As indicated earlier in **Figure 4**, the STDP-RL model performance did not only depend on the model seed but also the game initial state. Altogether the EVOL model showed optimal performance independent of the model and game initializations, thereby also demonstrating robustness.

**Figure 6:**
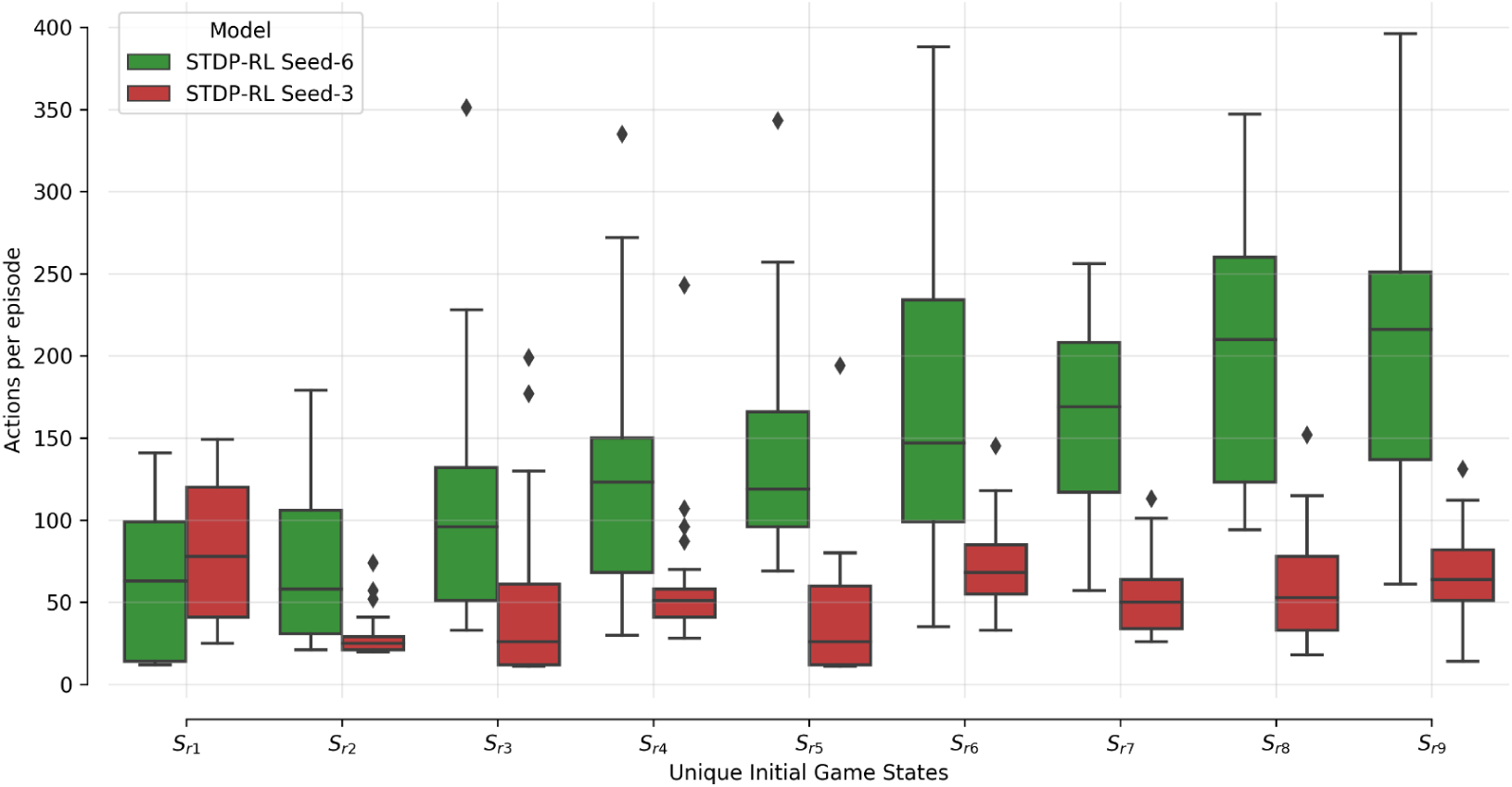
SNN STDP-RL trained models on different episodes. Simulations for nine different episodes (Sr1-Sr9) were repeated 25 times each, using STDP-RL Seed-6 (best-performing) and Seed-3 (worst-performing) models. Note that each episode had a unique initial game-state which was identical across repeats of the same episode.

### Individual neurons develop preference for a ‘specific’ action generation after training

Comparing the activity patterns of the models while playing CartPole may not reveal developed features of the full neuronal network possibly because of a different set of game-states evoked during each episode of CartPole. Therefore, to uncover the response properties of all network model elements, we simulated the trained models by sequentially activating all possible sensory inputs i.e. 20^4^ unique quadruplets (20 neurons to encode each of the 4 sensory parameters) and recorded the responses of all neurons as well as the associated action (i.e. if firing rate of EM-L > firing rate of EM-R, action will be Move-Left and if firing rate of EM-R >firing rate of EM-L, action will be Move-Right) in the SNN model. During these activations, we kept the model isolated from the game, to allow full control of the model’s sensory state.

Activation of each sensory neuron led to the activation of many neurons at multiple postsynaptic levels, pushing the model towards making a decision (Move-Left or -Right). Since the decisions were always based on the population level activity of motor areas, it was nontrivial to establish causality. Instead, we compared the average action selected when each sensory neuron is activated and the cascading activations throughout the rest of the network (**Figure 7**). For each activated sensory neuron (y-axis in **Figure 7**), we first counted the number of times each neuron of the network (x-axis in **Figure 7**) triggered during a ‘Move-Left’ or ‘Move-Right’ action. Further, we assigned it a Move-Left or Move-Right preference (heatmap color in **Figure 7**) if it was more active during Left or Right actions, respectively. The colormaps (**Figure 7**), show the normalized count of preferred action generation contribution of each neuron in the network referred to as ‘Action selectivity’ later in the text. Note, we used -ive sign with ‘Action selectivity’ to indicate ‘Move-Right’ actions and +ive sign to indicate ‘Move-Left’ actions. Before training (**Figure 7A**), all the neurons in the circuit were minimally biased towards each action as ‘Action selectivity’ values shown in the colormap are 0 (stay), marginally greater than 0.5 (for ‘Move-Left’ action), or smaller than -0.5 (for ‘Move-Right’ action). After training (**Figures 7B-D**), the preference of some neurons increased towards a single action as the range of colors in the map expanded towards +1 (for left action) and -1(for right action).

**Figure 7:**
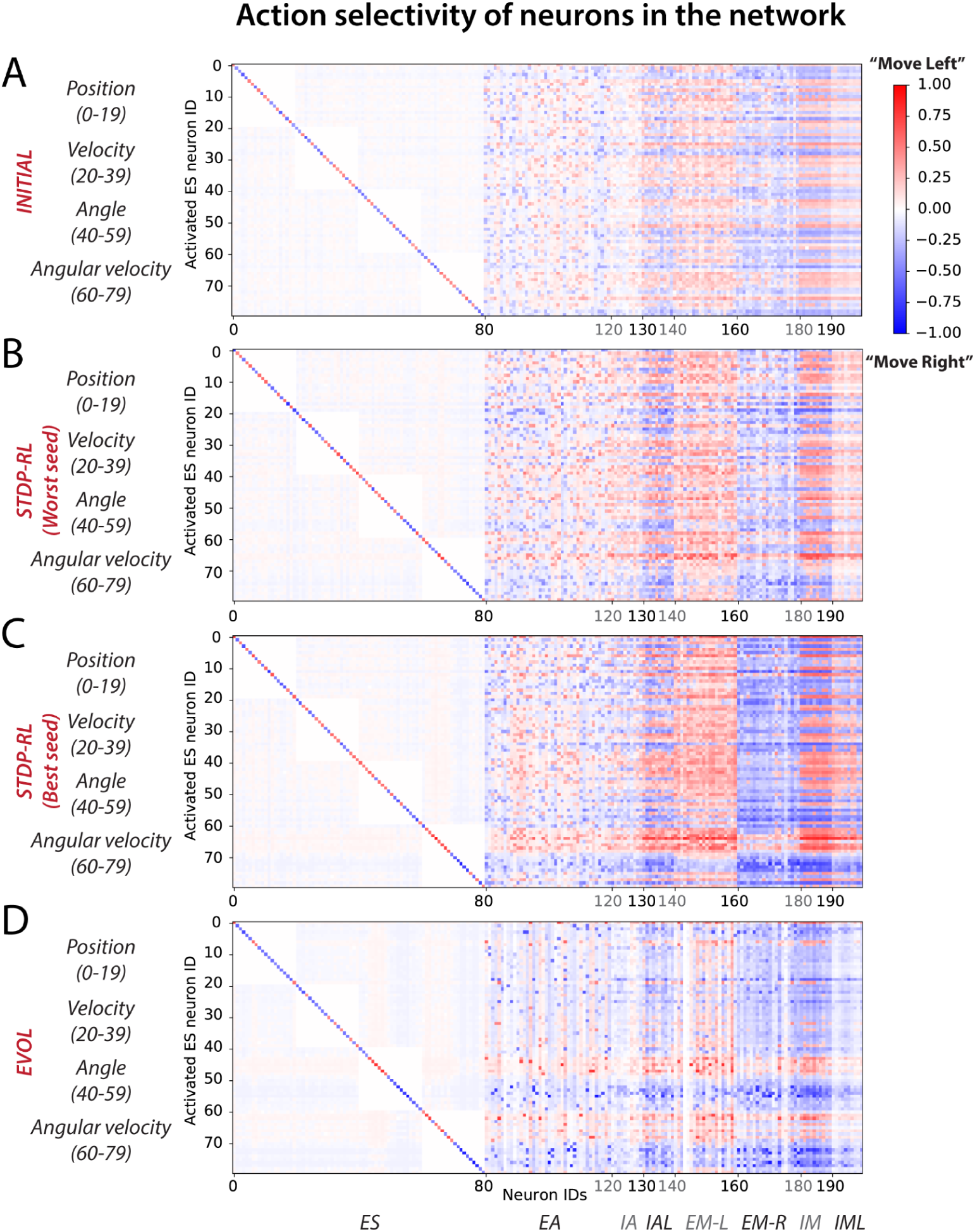
Action selectivity maps show differential participation of neurons in the network in action generation. In response to a particular sensory input, each neuron was considered to contribute to Move-left or Move-right actions and then labeled with the frequency of the number of times it was activated relative to the frequency of those actions (with positive sign indicating activations relative to number of moves left, and negative sign indicating activations relative to moves right). **A)** INITIAL, **B)** STDP-RL (Worst seed), **C)** STDP-RL (Best seed), and **D)** EVOL. Note that propensity towards an action is defined as the number of times a neuron is active during a particular action choice. For example, if out of 8000 sensory inputs, 6000 times the action was Move left and 2000 times the action was Move Right, for each of those sets of conditions we compute how many times each neuron was active. If the neuron was active 3000 out of the 6000 moves left, then the participation value is 0.5. If the same neuron gets activated 2000 times to move right, then its participation value is -1 (negative sign is for move right, positive for move left).

In the STDP-RL model with the worst performance (Worst Seed in **Figure 7B**), most of the neurons in EM-L (EM-R) participated dominantly in generating ‘Move-Left’ (‘Move-Right’) actions without any clear preference for particular input features, similar to the INITIAL network (**Figure 7A**). For sparser input values of ‘Position’ and ‘Angular Velocity’, some neurons in EM-L flipped their preference slightly to ‘Move Right’ (blue colored pixels for EM-L neurons in **Figure 7B**), and some neurons in EM-R flipped their preference slightly to ‘Move Left’ (red colored pixels for EM-R neurons in **Figure 7B**). Any effect on behavior due to sensory input parameters which were not used as ‘critic’ (e.g. ‘Position’ and ‘Velocity’) for training using STDP-RL were surprising. In contrast, in the STDP-RL model with the best performance (Best Seed in **Figure 7C**), most of the excitatory motor neurons in the circuit developed a non-overlapping association between positive angular velocity (ES neuron IDs 70-79) and ‘Move-Right’ action, and negative angular velocity (ES neuron IDs 60-69) and ‘Move-Left’ action. This means almost all of the neurons in EM-L were robustly activated only when the game-state had a negative angular velocity, and almost all of the neurons in EM-R were robustly activated only when the game-state had a positive angular velocity. We observed somewhat similar developed preferences for neurons in the EVOL model (**Figure 7D**), except that for positive or negative angular velocity, only a subset of EM-L and EM-R were activated i.e. not all EM-L or EM-R neurons were showing the same level of preference for the actions. In addition to angular velocity, we observed a similar effect for angle as neurons in EM-L were activated to generate ‘Move-Left’ actions only for positive angles as indicated by Neurons 40-50, whereas neurons in EM-R were activated to generate ‘Move-Right’ actions only for negative angles. The emerging preference of EM-L and EM-R neurons for positive and negative values of angles and angular velocities in EVOL model (**Figure 7D**) might explain its superior performance over models trained with STDP-RL where we observed such preference only for angular velocity (**Figure 7C**). Moreover, despite being not used for training, an emerging preference for ‘Angle’ and ‘Angular velocity’ in the SNN trained using EVOL showed that these two parameters were sufficient to capture sensory-motor behavior tested in this study.

For the EVOL model we also observed a complete deactivation of some neurons (vertical white lines in **Figure 7D**) that might have been counterproductive in performing a correct movement. The EVOL model showed a greater variability in the final total reception weight for postsynaptic neurons, including greater weight decreases that effectively deactivated neurons (**Supplementary Figure 4**). This neuron pruning strategy was not employed by the STDP-RL learning algorithm and might be a potential reason for the stunted game performance and the variability based on initial connections.

### Training using EVOL enabled SNN to learn broader sensory-motor associations

Plotting the activity levels of neurons during specific actions in response to each pair of sensory-inputs could diffuse the learned sensory-motor associations, especially since the model learned not about individual sensory parameters but a full game-state consisting of four sensory parameters at each game-step. Therefore, we next computed sensory-motor maps for pairs of sensory input parameters (**Figure 8**). Before training, for only a few pairs of sensory input parameters, the model had weak preference either to move left, right, or stay (**Figure 8A**). After training using STDP-RL, the model with worst performance (marked with Worst Seed in **Figure 8B**) developed a modest action preference for a few values of position (see the horizontal yellow line at *Position: “18”* for ***“Move Right”*** in **Figure 8B**), velocity (see horizontal yellow lines at *Velocity*: *“14”* and *“16”* for ***“Move Right”***,*“13”* and *“15”* for ***“Move Left”*** heatmaps in **Figure 8B**) and angles (see horizontal yellow lines at *“4”, “8”* and *“14-17”* for ***“Move Right”*** *and “7”, “9”, “13” and “18”* for ***“Move Left”*** heatmaps in **Figure 8B**) but somewhat stronger action preference for positive and negative values of *Angular velocity* around *“5 and 6”* for ***“Move Right”*** and *“14”* for ***“Move Left”*** heatmaps (angular velocity; **Figure 8B**). On other hand, the model with best performance (marked with Best Seed in **Figure 8C**) developed a stronger action preference for a broader range of positive and negative values of Angular velocity around “10” for “Move Right” and “Move Left” heatmaps (see broader yellow bands on *“0-9”* for ***“Move Left”*** and *“10-19”* for ***“Move Right”*** in **Figure 8C**) in addition to some input values of other sensory parameters. These results are in line with results in **Figures 7B-C**, confirming that the STDP-RL trained models only learned robust action preference based on Angular velocity and could not learn from any other parameter robustly. In contrast, in models using EVOL, we found quite a few input-output associations (weak association for *Position*: *“0”, “2”, “6-7”* and *“19”* for ***“Move Left”*** and *“1-5”, “8-18”* for ***“Move Right”*** heatmaps; stronger association for *Angle*: *“0-9”* for ***“Move Left”*** and *“10-19”* for ***“Move Right”*** heatmaps; stronger association for *Angular velocity*: *“0-9”* for ***“Move Left”*** and *“10-19”* for ***“Move Right”*** heatmaps in **Figure 8D**), indicating distributed learned associations between actions and some values of all sensory input parameters. These results clearly indicate that better performance of models trained using EVOL could be attributed to use of more sensory information in decision making because the model learned to take action based on two sensory signals i.e. Angle and Angular velocity, in contrast with STDP-RL trained models, which learned to take action only based on Angular velocity. However, some of these differences could be due to the fact that the STDP-RL critic explicitly utilized two parameters: angle and angular velocity (see Methods). Other critics for STDP-RL could produce different outcomes.

**Figure 8:**
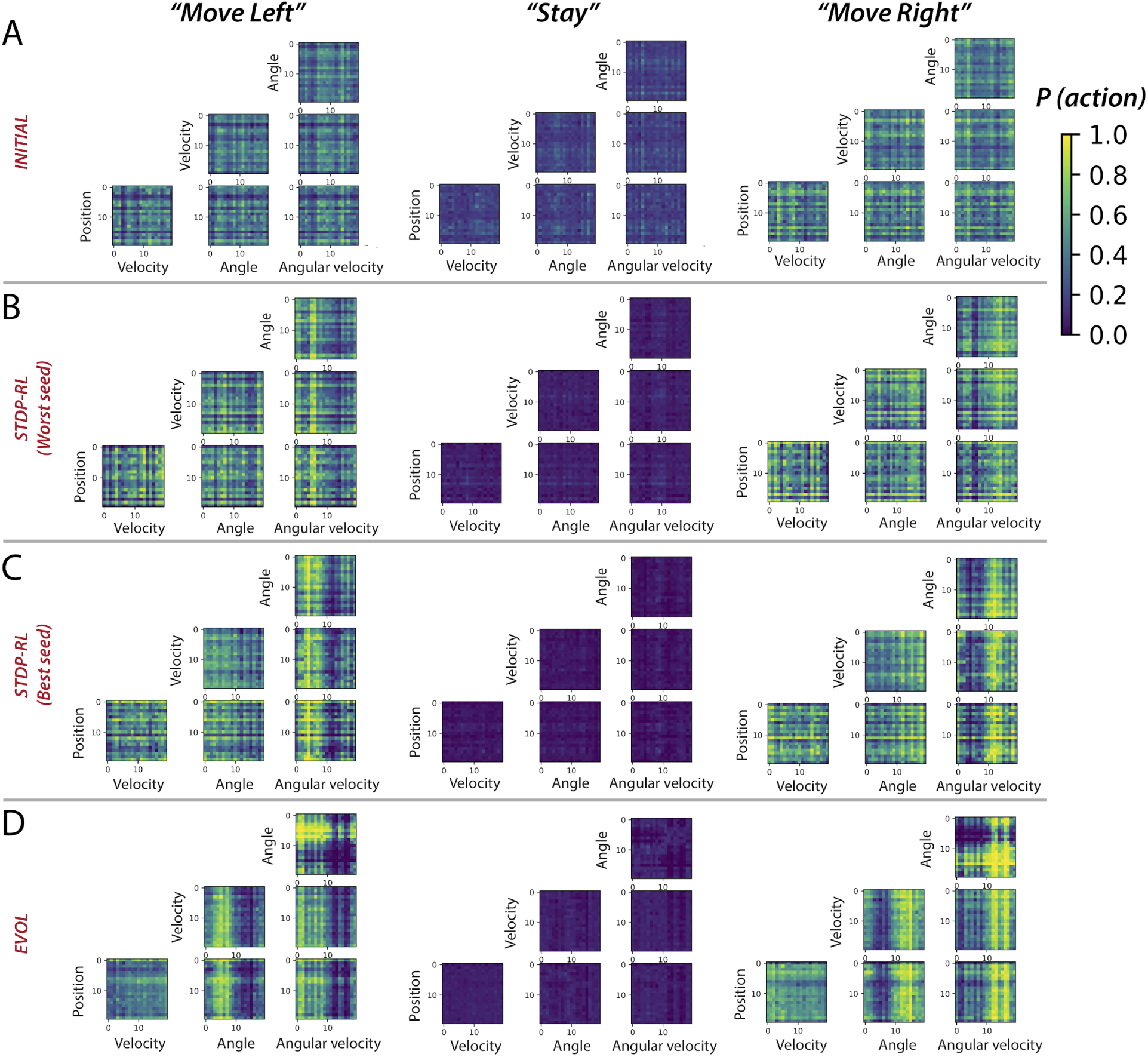
Sensory-Motor mappings show different behaviorally-relevant input-action associations learned using STDP-RL and EVOL strategies. Probability of “Move Left” (left panel), “Stay” and “Move Right” for all pairs of sensory input parameters for the model before training **(A)**, after training using STDP-RL **(Worst seed: B)**, after training using STDP-RL **(Best seed: C)**, and after training using EVOL **(D)**.

## Discussion

In this work, we developed and trained an SNN using biologically inspired STDP-RL and evolutionary strategy (EVOL) algorithms. One of our goals was to investigate biologically-plausible learning algorithms that operate at different timescales, and determine their strengths and weaknesses, to enable offering insights into biological processes. Comparing the performance of our SNN model trained using STDP-RL and EVOL (**Figure 4 vs Figure 5 and Table 3**), we demonstrated that both strategies could be used for training our SNNs to play CartPole. There were also noticeable differences in resulting neuronal dynamics and behavior (**Figures 7-8**). The evolutionary strategy showed excellent performance training our SNNs to play CartPole (**Figure 5**).

As mentioned, the design of the two algorithms is substantially different in several regards. First, STDP-RL relies on an explicit hand-crafted critic, which in our case used two variables (angle, angular velocity), to inform the model of the quality of its decisions. Second, the STDP-RL critic used short intervals to evaluate the model, close to the time-scale of individual moves. In contrast, EVOL evaluated the quality of a model using only the game score, evaluated over substantially longer episodic time-scales. Some of these differences are not intrinsic to the individual algorithms. However, EVOL allows a more flexible design, and the longer time-scales used potentially offer more accurate evaluation of a model’s quality.

It is interesting to see how training our models using these different strategies produced similar sensory-motor associations without explicitly training the models to execute specific motor actions for a given game-state. Regardless of the training algorithm, all models learned to associate actions dominantly to specific values of *Angular velocity* (**Figures 7-8**). Analyzing the single input-action mappings (**Figure 7**), we did not find clearly strong or broad associations between the Position, Angle, and Velocity with actions, which appeared in analyzing the pairwise input-action mappings (**Figure 8**). It is possible that the associations between sensory inputs and motor actions for other parameters were not fully revealed because the SNN was trained using all four parameter values, and some weak associations between parameters other than Angular velocity may be broadly present, but did not appear in analysis which was limited to pairwise inputs. As noted above, the STDP-RL critic also depended upon only two parameter values, potentially explaining these findings. Another reason for limited tuning for some parameters could be that during training the receptive fields for those parameters were not experienced by the model, and therefore the model could not learn any specific associations for those inputs. This is indicated in the receptive fields parsed during the episodes of the game played after training (**Figure 7**).

Our modeling underlines the benefits to training from a set of multiple initial network configurations, achieved through varying synaptic connection weights or network architectures (Stanley and Miikkulainen 2002; Neymotin et al. 2013), and testing the populations’ performance. The best performing model can then be used for longer-term training. An alternative is to use multiple models from the population, and ideally promote model diversity. We highlight these issues by means of the training strategies used in this work. In our STDP-RL model, we first selected a set of hyperparameters suitable for optimizing learning and then tuned for initial synaptic weights using the fixed optimal hyperparameter values. Thus, the initial weights and the learning parameter values differed from those at the beginning of the hyperparameter search. Furthermore, when we trained our SNN model using STDP-RL and a different set of initial weights (by changing random numbers), we observed variable performance, showing large influence of initial parameters of learning behavior. In our EVOL strategy, without using any explicit synaptic learning mechanism, we evaluated the model’s performance with different synaptic weights and found synaptic weight distributions showing good performance. Randomizing initial weights did not impact the performance as strongly as in STDP-RL trained models. Comparing the performance of models using these two training strategies (**Figure 3** vs **Figure 5**), we clearly showed that EVOL produced models with robust and optimal performance. Although models trained using EVOL generated different distributions of synaptic weights, the robust peak performance was demonstrated, which is consistent with the observed variability in circuit elements of biological neuronal circuits (Marder and Goaillard 2006; Calabrese et al. 2011; Bucher, Prinz, and Marder 2005; Calabrese, Norris, and Wenning 2016; Marder and Taylor 2011; Marder 2011; Roffman, Norris, and Calabrese 2012; Golowasch 2014; Hamood and Marder 2014; Goaillard et al. 2009; Anwar et al. 2022).

In the past, the use of evolutionary algorithms in neurobiological models has mainly been limited to optimizing individual neurons (Werner Van Geit, Achard, and De Schutter 2007; W. Van Geit, De Schutter, and Achard 2008; Rumbell et al. 2016; Neymotin et al. 2017), or neuronal networks through hyperparameter tuning (S. Dura-Bernal et al. 2017). Although more recent work makes changes to network architectures (Stanley et al. 2019), modifications of synaptic weight matrices in spiking or biophysical neuronal networks have rarely been performed, partly due to the large computational costs associated with searching through the high dimensional space. Here we have demonstrated that evolutionary algorithms operating on synaptic weight matrices are an effective strategy to train SNNs to perform behaviors in a dynamic environment.

We previously used the STDP-RL learning rules to train a visual/motor cortex model to play Pong (Anwar et al. 2021). That model required additional complexity for encoding the visual scene (object location, motion direction). This complexity made it more challenging to decipher the role of different components/parameters of the learning algorithms and how to optimize them. In the present SNN model, the lack of visual cortex was a simplification that allowed us to perform a more extensive hyperparameter search to increase the chances for STDP-RL to succeed. After hyperparameter optimization, the STDP-RL algorithm was effective in producing reasonable performance in CartPole. Furthermore, the use of CartPole and the simpler sensory/motor cortex model also allowed us to test long optimizations using EVOL in parallel on supercomputers. In the future, with the knowledge gained here, we will test our new algorithms using more complex models, tasks, and environments.

Despite the substantial differences between ANNs and SNNs, learning in both is primarily realized by adjusting the weights of connections or synaptic strengths among interconnected neurons. In ANNs, this usually occurs through the following sequence, repeated many times: 1) inputs are sampled, 2) corresponding outputs are evaluated, and 3) weights are adjusted to minimize the output error via back-propagation through hidden network layers (Schmidhuber 2015). In SNNs, the weights are often adjusted using hebbian or spike-timing dependent plasticity (STDP) rules (Dan and Poo 2004, 2006; Izhikevich 2007; Farries and Fairhall 2007; Caporale and Dan 2008). These strategies are useful when there is a temporally proximate relationship between inputs and outputs and there is no feedback involved.

Sensory-motor RL is a more difficult problem to solve because behaviors require evaluation of many sub-actions, and are associated with different environmental cues integrated over time. Moreover, the reward/punishment feedback is delivered later, which makes it difficult to attribute reward/punishment only to relevant actions and neuron groups producing those actions (Izhikevich 2007). Recent ANNs have taken advantage of a *replay and update* strategy to re-sample previous experiences and shape the action policy for given sensory cues maximizing the cumulative reward (Hayes et al. 2021). In SNNs using STDP-RL, eligibility traces can be used to associate reward/punishment to corresponding actions and neuronal assemblies backwards in time, as we have implemented in our work here: when a postsynaptic neuron fires within a few milliseconds of presynaptic neurons firing, a synaptic eligibility trace is activated, allowing the synapse to undergo potentiation or depression during the following 1-5 seconds (Anwar et al. 2021; Patel et al. 2019; Hazan et al. 2018; Chadderdon et al. 2012). STDP-RL trains SNNs by establishing associations between the neurons encoding the sensory environment and neurons producing actions or sequence of actions, such that appropriate actions are produced for specific sensory cues. The sensory-motor associations are established from reward-modulated synaptic weight changes that occur at each timestep of the simulation. STDP-RL trains at the individual model level, as we consider each separate initialization of an SNN network a separate “individual”.

In contrast to individual-based algorithms such as STDP-RL, evolutionary algorithms operate at vastly different timescales, and typically use populations of models (Feldman, Aoki, and Kumm 1996; Parisi et al. 2019). Evolution is successful when individuals who are fit enough to produce offspring pass their genes to the next generation (Garrett 2012). While individual learning is restricted to an animal’s lifespan, it still confers powerful competitive advantages. This, in turn, feeds into the evolutionary process: animals that learn the idiosyncrasies of their environment, including its threats and rewards, are more likely to survive and propagate. The obvious genomic storage limitations prevent the encoding of all important environmental information within an animal’s genome (Zador 2019; Koulakov, Shuvaev, and Zador 2021), underlining how individual learning must complement evolution. As noted by James Mark Baldwin, learning on an individual level drives the evolutionary process (“A NEW FACTOR IN EVOLUTION” 1896). To simulate this process, we envision a strategy that implicitly choses the next generation’s weights and connectivity patterns based on learning success, thereby accelerating population-level fitness improvements. This is consistent with the accelerated learning of models when deploying the Baldwin effect, as has been recently demonstrated in embodied intelligence tasks (Gupta et al. 2021). The extent to which these strategies translate to more complex tasks and circuit architectures could offer further insights into multi-scale neurobiological learning.

## Acknowledgments

This research was funded by Army Research Office W911NF-19-1-0402, Army Research Office Undergraduate Research Apprenticeship Program supplement, Army Research Lab Cooperative Agreement W911NF-22-2-0139, and National Institute on Deafness and Other Communication Disorders R01DC012947-06A1. The authors acknowledge the Tufts University High Performance Compute Cluster (https://it.tufts.edu/high-performance-computing) which was utilized for the research reported in this paper. In addition, some of the calculations done in this work used the Extreme Science and Engineering Discovery Environment (XSEDE) Bridges GPU Artificial Intelligence at Pittsburgh Supercomputing Center through allocation CCR200032. The views and conclusions contained in this document are those of the authors and should not be interpreted as representing the official policies, either expressed or implied, of the Army Research Office or the U.S. Government. The U.S.Government is authorized to reproduce and distribute reprints for Government purposes, notwithstanding any copyright notation herein.

## Contribution to the field statement

Biological learning operates at multiple timescales, over long evolutionary stretches, down to the learning that takes place during an individual’s lifetime. While each process has been simulated as a basic learning algorithm in the context of spiking neuronal networks (SNNs), investigations and comparisons of the two have remained limited. In this study, we train SNNs to play the CartPole game using: 1) learning during a model’s lifetime through spike-timing dependent reinforcement learning (RL) and 2) simulated evolutionary learning processes. We evaluate the performance of each algorithm after training and through the creation of sensory/motor action maps that delineate the network’s transformation of sensory inputs into higher-order representations and eventual motor decisions. While both algorithms produced SNNs capable of moving the cart left and right and keeping the pole vertical, evolutionary algorithms displayed superior performance, and are therefore a more powerful method to train SNNs to perform sensory-motor behaviors. Our modeling opens up new capabilities for SNNs in reinforcement learning tasks and could serve as a testbed for neurobiologists aiming to understand the multiple timescales of learning and dynamics in the brain.

## Supplementary Figures

**Supplementary Figure 1:**
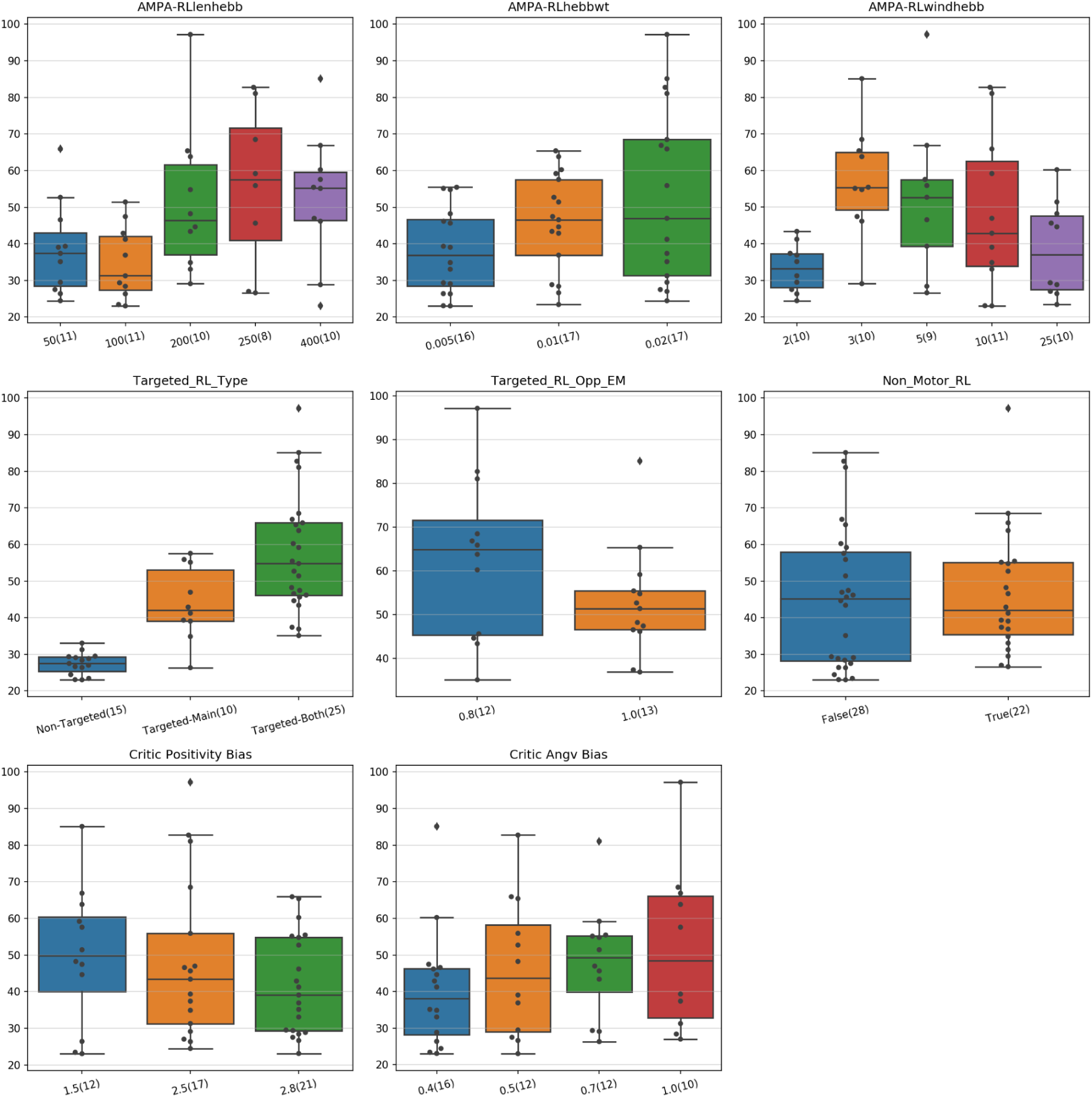
Performance distribution of the first hyperparameter search for training using STDP-RL (displaying averages over 100 episodes during training). In each panel, the y-axis shows the performance, and the x-axis indicates the parameter values used in the evaluation. The number of models run with the specific parameter value is presented in parentheses, with 107 tested hyperparameter combinations.

**Supplementary Figure 2:**
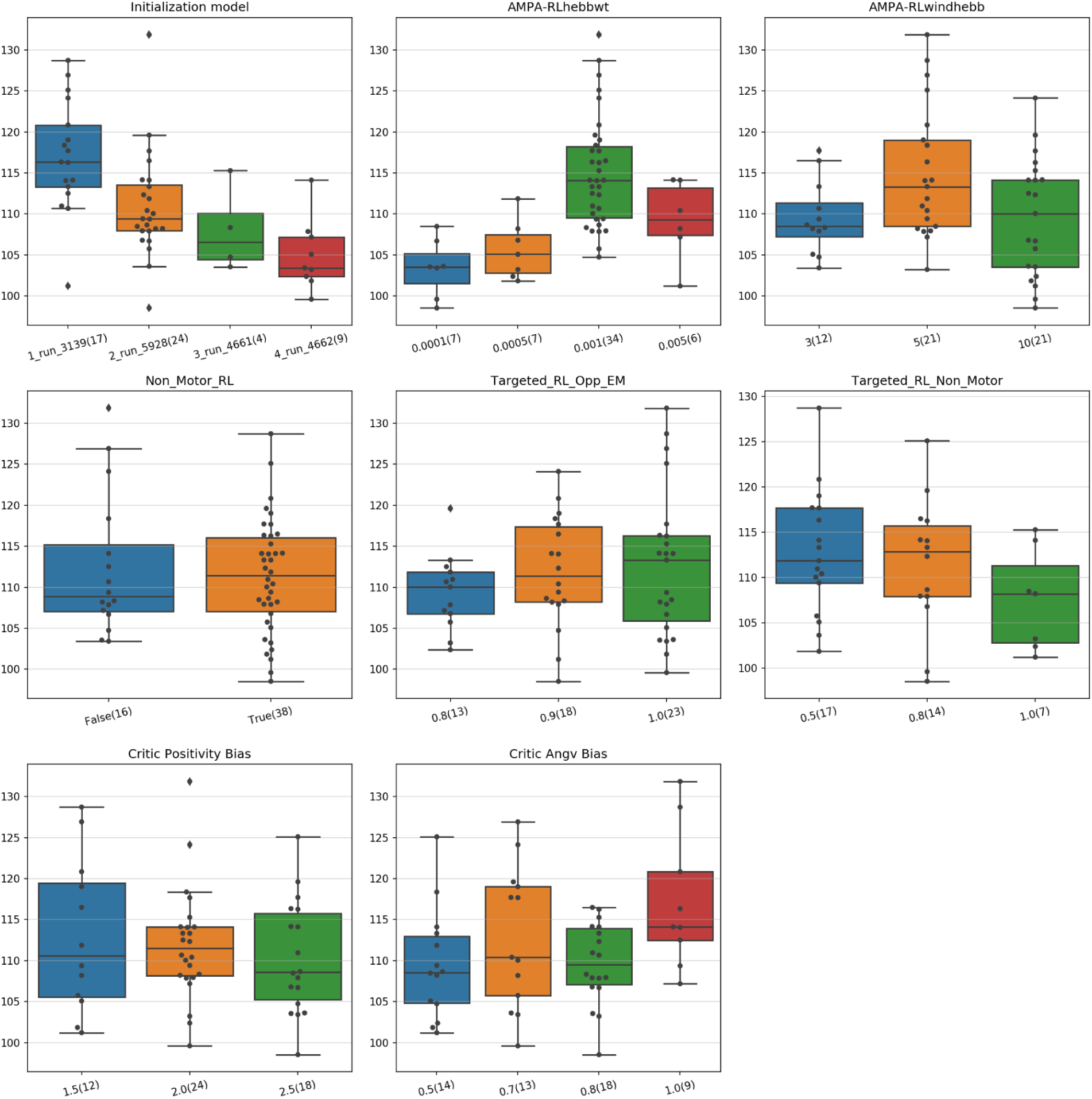
Performance distribution of hyperparameters for training using STDP-RL evaluated in the second step (using averages over 100 episodes).

**Supplementary Figure 3:**
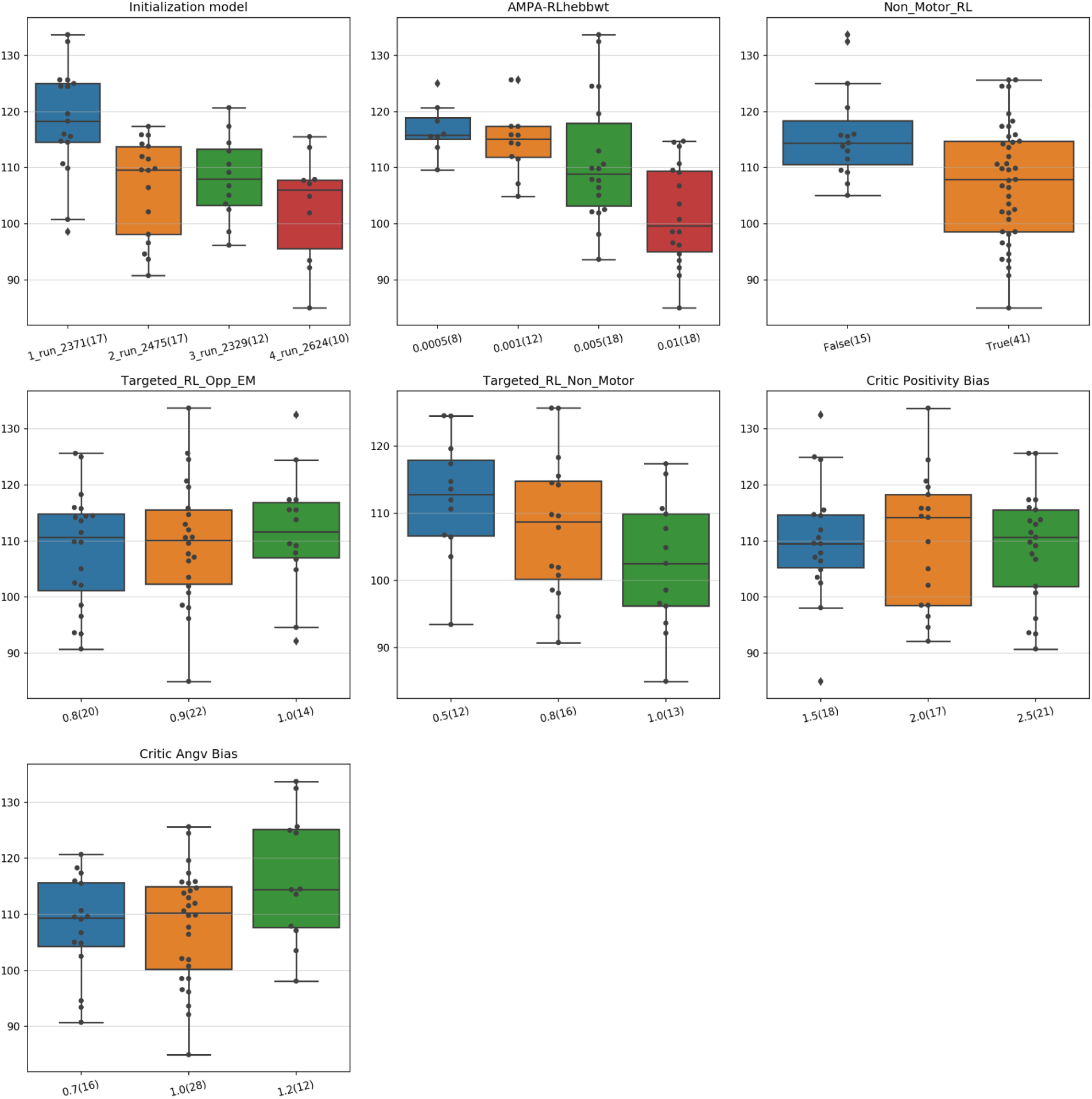
Performance distribution of hyperparameters for training using STDP-RL evaluated in the third step (using averages over 100 episodes).

**Supplementary Figure 4:**
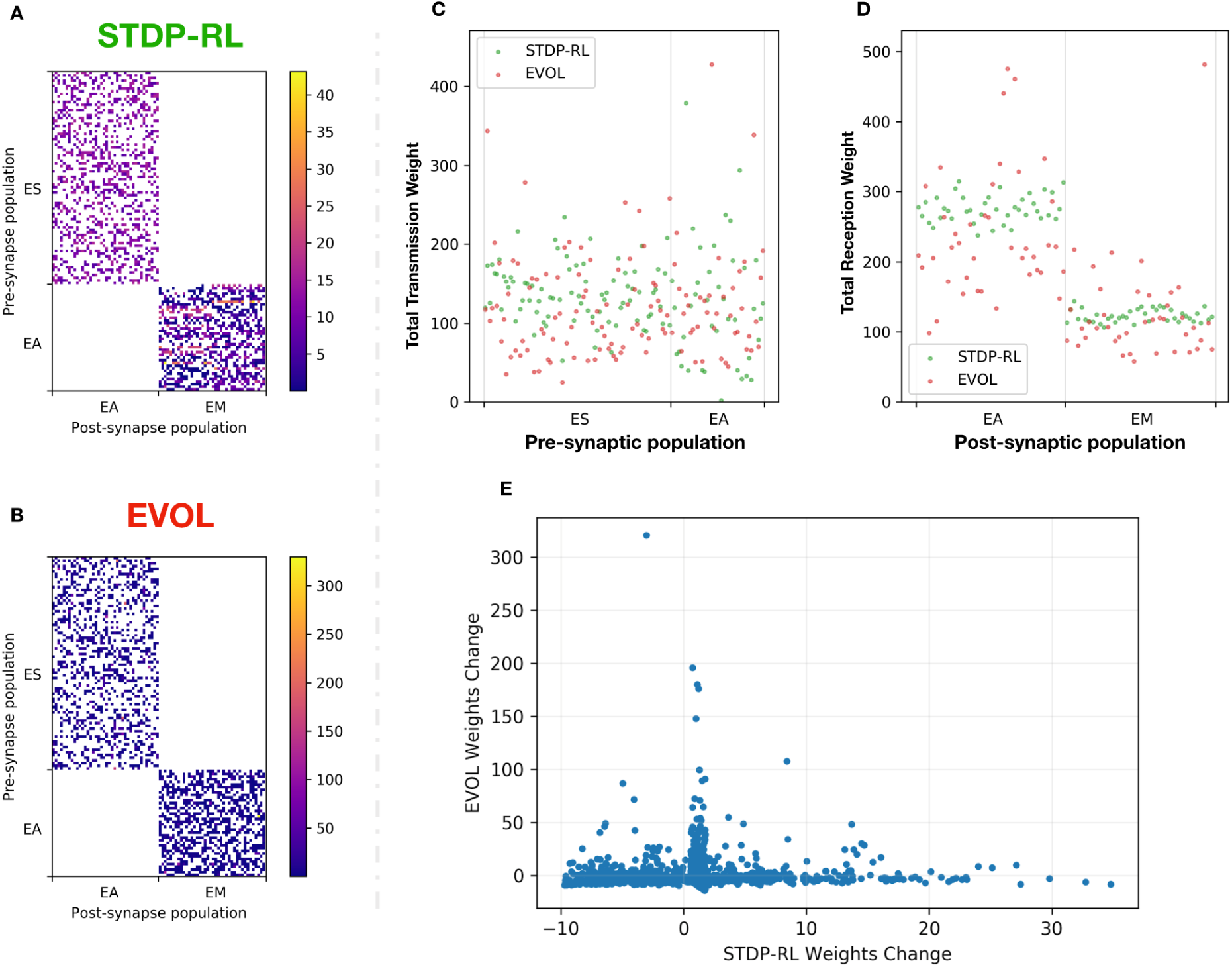
STDP-RL and EVOL differentially modulate the synaptic weights of the SNN model during training. **(A-B)** Adjacency matrices showing the weights of synaptic connections between ES-EA and EA-EM populations in STDP-RL model **(A)**, EVOL model **(B)**. Note that the adjacency matrices have different scales as EVOL model has weights reaching 300. **(C)** Total Transmission Weight: Sum of synaptic weights from a presynaptic neuron onto multiple postsynaptic neurons. **(D)** Total Reception Weight: Sum of synaptic weights onto each postsynaptic neuron from multiple presynaptic neurons. **(E)** As each model changed the weights from the original initialization, each dot represents a synaptic connection weight after training with STDP-RL (x-axis) and EVOL (y-axis) with a Spearman correlation of -0.09 (p-value = 0.0001).

**Supplementary Table 1:** STDP-RL parameters obtained through hyperparameter search.

| Parameter | Hyperparameter Search Step 1 | Hyperparameter Search Step 2 | Hyperparameter Search Step 3 | Description |
| --- | --- | --- | --- | --- |
| Duration (s)<br>(not varying) | **500** | **2000** | **2000+** | Duration in seconds that was trained using each. A step was performed in 50ms |
| AMPA-RLwindhebb (ms) | (2, **3**, 5, 10, 25) | (3, **5**, 10) | **5** | Maximum time between presynaptic and postsynaptic spike times for considering plasticity. |
| AMPA-RLlenhebb (ms) | (50, 100, 200, 250, **400**) | **250** | **250** | The decay time constant of the exponentially decreasing eligibility trace. |
| AMPA-RLhebbwt | (0.005, 0.01, **0.02**) | (0.0001, 0.0005, **0.001**, 0.005) | (0.0005, 0.001, **0.005**, 0.01) | Max synaptic weight adjustments based on reward or punishing signal. |
| Targeted_RL_Type | (non-targeted RL, targeted RL main, **targeted RL both**) | **targeted RL both** | **targeted RL both** | The Targeted RL paradigm chosen |
| Non_Motor_RL | (**False**, True) | (**False**, True) | (**False**, True) | Activating STDP-RL plasticity for synapses in the $ES \rightarrow EA$ pathway |
| Targeted_RL_Opp_EM | (0.8, **1.0**) | (0.8, 0.9, **1.0**) | (0.8, **0.9**, 1.0) | Attenuation factor for the opposite subpopulation receiving reinforcement |
| Targeted_RL_Non_Motor | **1.0** | (0.5, 0.8, **1.0**) | (0.5, 0.8, **1.0**) | Attenuation factor for the non motor subpopulation receiving reinforcement |
| Critic Positivity Bias | (**1.5**, 2.5, 2.8) | (1.5, **2.0**, 2.5) | (1.5, **2.0**, 2.5) | $\eta_{angvel}$: used in critic (**equation 1**) |
| Critic Angv Bias | (**0.4**, 0.5, 0.7, 1.0) | (0.5, 0.7, 0.8, **1.0**) | (0.7, 1.0, **1.2**) | $\eta_{positivity}$: defined for critic evaluation (**equations 2-3**) |

